# Putting cognitive tasks on trial: A measure of reliability convergence

**DOI:** 10.1101/2023.07.03.547563

**Authors:** Jan Kadlec, Catherine Walsh, Uri Sadé, Ariel Amir, Jesse Rissman, Michal Ramot

## Abstract

The surge in interest in individual differences has coincided with the latest replication crisis centered around brain-wide association studies of brain-behavior correlations. Yet the reliability of the measures we use in cognitive neuroscience, a crucial component of this brain-behavior relationship, is often assumed but not directly tested. Here, we evaluate the reliability of different cognitive tasks on a large dataset of over 250 participants, who each completed a multi-day task battery. We show how reliability improves as a function of number of trials, and describe the convergence of the reliability curves for the different tasks, allowing us to score tasks according to their suitability for studies of individual differences. To improve the accessibility of these findings, we designed a simple web-based tool that implements this function to calculate the convergence factor and predict the expected reliability for any given number of trials and participants, even based on limited pilot data.

## Introduction

One of the longstanding mysteries in cognitive science is what drives the profound variation in behavior that can be observed on almost any task, even within the typically developing population^1–4^. The search for the neural underpinnings of these individual differences, i.e., the neural variance between individuals that explains the behavioral variance, has been one of the key motivating forces behind cognitive neuroscience research in the last couple of decades^5–9^. Such brain-behavior association studies have demonstrated links between individual neural activation levels during tasks^10–12^ and behavioral performance, as well as between resting state brain networks and behavior^13–15^. This fascination with individual differences, which are found in animals as well as humans^16–18^, is driven not only by the deepest philosophical questions regarding individuality and our conception of the self but also by more practical considerations. Understanding individual differences will increase the utility of personalized biomarkers and treatments, which form the basis of precision medicine, and could revolutionize diagnosis and care for many disorders^19–22^. Yet this endeavor of linking brain and behavior, in particular through its occasionally problematic implementation in brain-wide association studies which carry out data-driven, whole brain searches for such correlations, has recently come under fire^23^. Criticism of the approach has centered on the small effect sizes, which can require thousands of participants in order to reveal a reproducible association.

Recent publications offer a variety of paths forward. One suggestion was that we must either settle for such very small effect sizes across very large populations (as in genome-wide association studies), or abandon the study of individual differences altogether, focusing instead on within-participant studies^24^. Others have highlighted the advantages of moving to more multivariate neural measures, especially when considering complex behaviors^25–27^, or the need to increase the reliability of the neural data^28,29^. Yet the effects of the reliability of the behavioral measurement, the oft-ignored second variable in this brain-behavior relationship, have garnered far less attention, although a few recent papers have begun to address this gap^30–37^.

Focusing on functional magnetic resonance imaging (fMRI) and taking a step back from this discussion, we must first consider what exactly it is we are measuring with brain-behavior correlations. In many cases, these correlations are calculated between a neural state and a behavior that are not measured simultaneously. This is always the case when the behavior is compared to resting state measures, but it is often also true when the neural state is derived from task-based measures^14,28,38,39^. Given the constraints of fMRI scanning, behavioral measures are usually collected outside the scanner, sometimes on different days than the scan itself. The underlying assumption for any such brain-behavior correlation to be meaningful must therefore be that we are measuring stable, trait-like behaviors.

Though it is generally implicitly assumed, behavioral tasks do not explicitly test whether the behaviors they are measuring indeed converge to a stable mean at the individual level. Test-retest reliability measures for most tasks are modest at best (∼0.6 reliability^40–48^). Is this due to a lack of convergence to a more stable performance, perhaps driven by momentary, non-specific fluctuations in a non-measured variable (attention, for instance), which are greater than the between-individual differences on the task? Or is this due to insufficient data being used to estimate individual abilities? In other words, will individual performance on a given task converge given a sufficient number of trials, or are these test-retest measures the true ceiling of reliability?

This question of convergence is crucial not only because reliability has an impact on the expected results of correlational analyses such as the relationship between brain and behavior, but also in itself. On the practical side, the rate at which performance at the individual level on a particular task converges has implications regarding the amount of data we must collect in order to reliably estimate trait-like abilities at the individual level. Beyond that however, there are also theoretical implications, regarding the nature of the ability we are attempting to measure, and the validity of the task for this purpose.

To address these questions, we collected data from more than 250 participants on a large battery of 14 cognitive tasks comprising 21 different behavioral measures, spanning multiple cognitive domains, allowing us to directly compare different tasks in the same individuals. The battery included 12 commonly used tasks, as well as two new tasks that we created. To increase the number of trials we collect for each task while minimizing the effect of learning across trials, we used multiple alternate forms of each task. For each behavioral measure, we tested the reliability of individual performance using a permutation-based split halves analysis. We find that while with enough trials all tasks tested will eventually converge to stable performance at the individual level, even across days, the rate of convergence varies substantially between tasks, with some tasks requiring an untenable number of trials to reach stability. Moreover, as has been previously described in the seminal work by Spearman and Brown^49,50^ and later elaborated on by others^33,51–54^, convergence across different tasks followed similar patterns which could be used to predict reliability. We reformulated the Spearman-Brown prophecy and defined a convergence coefficient *C* that allowed us to directly compare this convergence across different tasks in the same individuals. We further defined this convergence coefficient analytically using only the mean and variance of the distribution for binary measures (e.g., measures such as accuracy which comprised 15 out of 21 measures in our dataset). The analytical model was validated using both extensive simulations and real behavioral data. We also tested the model on increasingly smaller subsamples of both the simulated and the real-world data, to anticipate potential error arising from small sample sizes which are common in experimental settings. Limited numbers of trials per participant resulted in a systematic bias, which we were able to correct for by using the law of total variance. This correction allowed us to generate very accurate predictions of reliability even given only minimal data.

In order to improve the quality of behavioral data and behavioral tasks used by the community, we have developed an online tool based on this model, which can calculate reliability given data from any existing dataset including small pilot experiments (https://jankawis.github.io/reliability-web-app/). Our hope is that this will facilitate better planning and design of behavioral tasks by providing researchers an estimate of the number of trials necessary to reach any desired reliability level. This tool can be used to inform study design before committing to larger and more expensive studies.

## Results

### Measuring reliability

In the study of individual differences, it is very common to calculate correlations between two different measures across participants. For instance, in brain-behavior correlations, the main methodology is to search for correlations between the neural variance across participants and the behavioral variance across those same participants. For this metric to be meaningful, both the neural and the behavioral data need to be reliable in the sense that participants’ ranking within the cohort is as consistent as possible, as it is this internal consistency which drives the brain-behavior correlations. This is also the case when we seek to measure correlations of individuals across different behavioral tasks, or the stability and internal consistency of any measure in general. Throughout this paper, we therefore define reliability (*R*) as the Pearson’s correlation across participants on different subsets of data from a given behavioral measure, using a permutation-based split-half extension of the classic test-retest reliability measure (see Methods)^51,52,55–57^. This method has previously been suggested to be optimal for calculating the internal consistency of cognitive tasks^33,51,57–60^. In order to calculate reliability for different sample sizes, we split participants’ data into two smaller, non-overlapping samples of size (*L*), calculate the scores from those samples in each split per participant, and correlate these two measures across participants using Pearson’s correlation. This correlation is an indication of the consistency of the relationship between participants on this task when averaging across *L* trials to estimate performance per participant. We then repeat this calculation 1000 times to eliminate noise coming from a specific split selection, giving us a distribution of reliabilities for this behavioral measure at this sample size. The mean of this distribution, which is equivalent to Cronbach’s alpha^55,56,61^, is used to create reliability vs number of trials (*RL*) curves for each task (Fig. 1). Those curves are theoretically bound between -1 (perfect negative correlation) and 1 (perfect correlation), where a reliability of 0 describes a random dataset with no internal consistency in the order of participants’ scores across different subsamples of data (for instance, a participant might be in the 93rd percentile of participants in one subset, and in the 62nd percentile in a different subset), while a reliability of 1 denotes a perfectly reliable dataset where the relationship between participants is consistently maintained across all data subsets.

**Figure 1.**
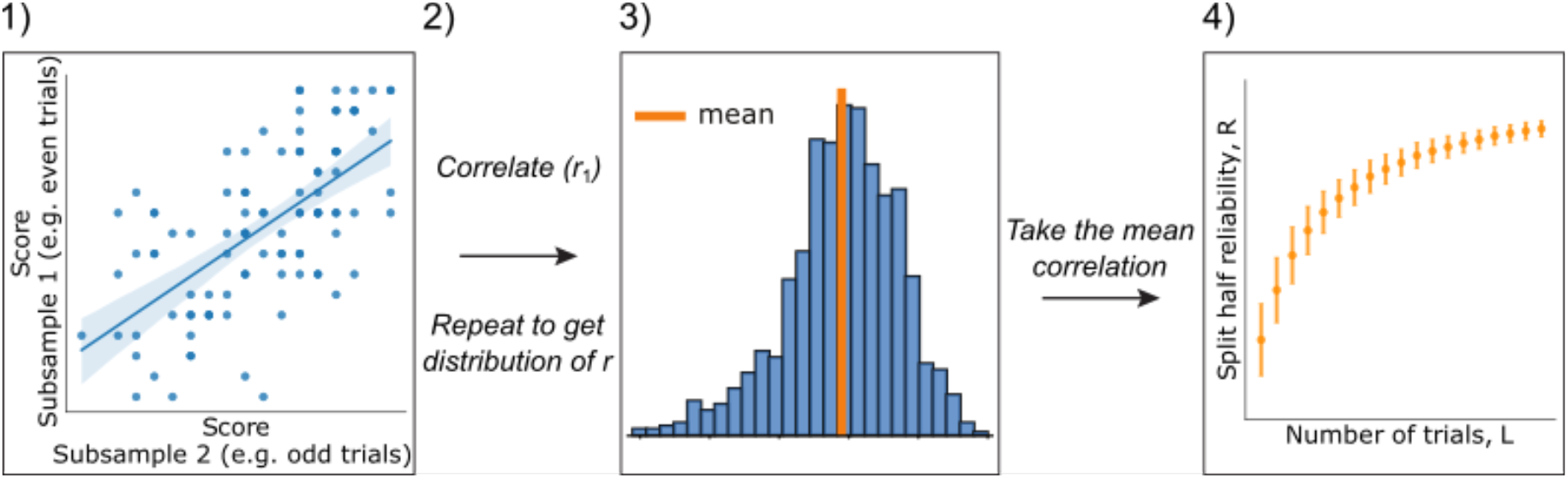
Reliability calculation. 1) For each of *N* participants, we take two random samples of trials L (up to a maximum of half the total trials in a task) and then calculate the score for each sample, creating two vectors of length N. We compute the Pearson correlation of these vectors of subsampled scores across participants to get the correlation, *r*_*1*_. 2) We repeat this process 1000 times to get a distribution of correlations. 3) We take the mean of this distribution, *R*, to create one point in the *RL* plot. 4) We repeat for other values of *L* to create the full *RL* curve, errors are standard deviations of the distribution of reliabilities.

We began by testing the reliability of a large battery of 12 commonly used cognitive tasks, as well as two new tasks which we developed for the purpose of studying personal identity memory (PIM) and face memory/perception (FMP). These tasks span 21 different behavioral measures in total, with some tasks including multiple measures which capture different aspects of behavior (Table 1, Supplementary Fig. 1, Methods). In order to test how reliability behaves across a diverse set of measures, as opposed to simple accuracy calculations, we included several commonly used measures for these tasks. All data were collected online using the Prolific platform (https://www.prolific.co/), and were rigorously cleaned to detect and remove participants who were not paying adequate attention to the tasks (see Methods).

**Table 1.**
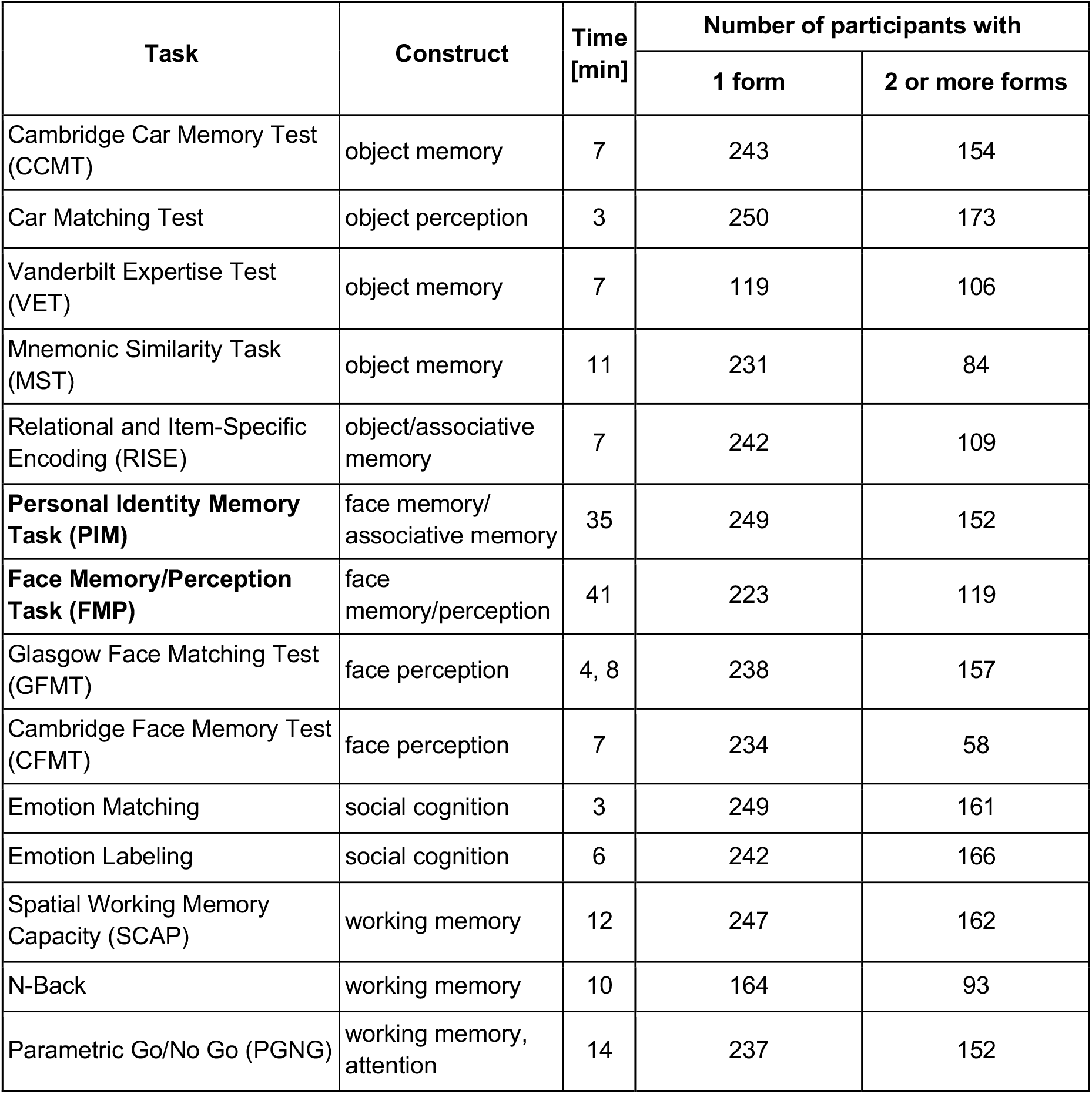
Table summarizing all tasks in our battery (first column) and the primary cognitive constructs (second column) that each is thought to estimate. Bolded items reflect the two custom tasks developed for this dataset. The third column details the mean time to complete a single form of this task (in the case of GFMT, it represents the time needed to complete the short and long versions of GFMT, respectively). The last two columns then detail how many participants completed at least one form of the task (left) or two or more forms (right). For tasks with two or more forms, this number indicates how many participants completed all available forms.

If the assumption of convergence to a stable mean holds, then averaging over more trials will provide a better estimate of participants’ true, stable trait-like proficiency, and our data will become more reliable. The rate of convergence is an indication of the degree of noise around each participant’s stable mean. Figure 2a-c shows examples of a reliable task, a less reliable task, and simulated random data, respectively (see Supplementary Figure 1 for reliability curves of all 21 measures). Note that for the random data, reliability converged to a spurious value, which across hundreds of simulations was bounded between -0.1 and 0.1, the average of which was not significantly different from zero (two-tailed t-test, t_99_=1.17, p=0.25). Small residual correlations in the random data might be a result of bias in the random seed used to generate the data, or of the limited sample size (see Methods)^62^. Data collection included at least twice as much data for almost all tasks than is commonly used (using alternate forms with different stimuli), allowing us to calculate the reliability of the number of trials contained in a standard one-form administration of the majority of our tasks (Fig. 2d). The initial data collection (one form of all tasks) was conducted over 3-4 days and took participants on average 193 minutes to complete. 183 subjects later completed additional forms of the tasks, consisting of an additional 185 minutes of data collection.

**Figure 2.**
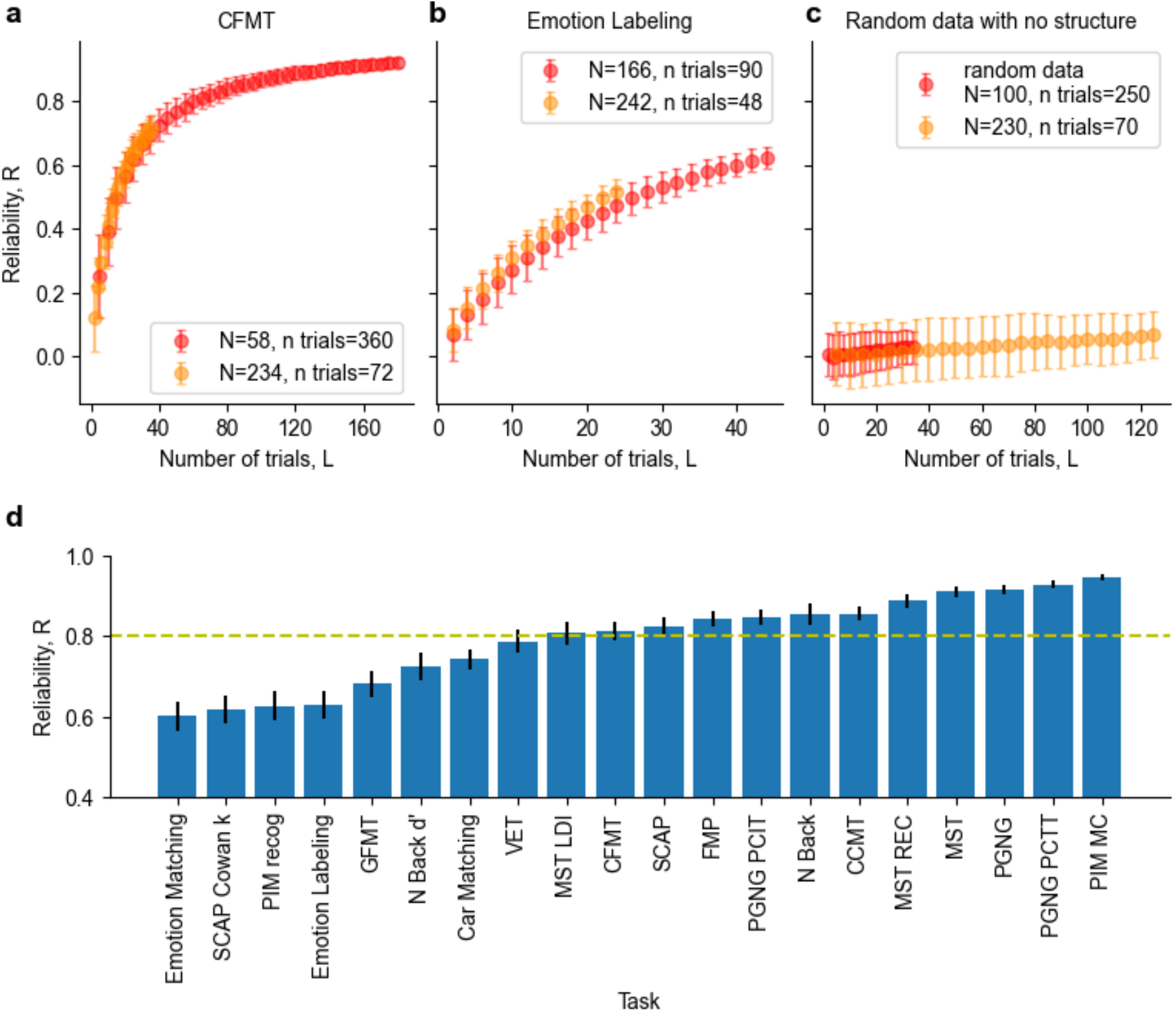
Dataset reliability. **a-c** *RL* reliability curves showing convergence for selected tasks as a function of number of trials. As fewer participants did all versions of the forms, we show here both samples with a larger *N* and fewer trials *L* (light orange), as well as smaller *N* with larger *L* (light red). **a**. Cambridge Face Memory Test (CFMT); **b**. Emotion Labeling task; **c**. randomly generated data showing no internal consistency. **d**. Barplot of reliabilities of one full form calculated from permutation based split-half reliability across all trials of two forms per measure. Dotted line represents 0.8 reliability. SCAP – Spatial Working Memory Capacity, PIM – Personal Identity Memory, GFMT – Glasgow Face Matching Task, VET – Vanderbilt Expertise Test, MST – Mnemonic Similarity Task, CFMT – Cambridge Face Memory Task, FMP – Face Memory Perception task, PGNG – Parametric Go-No Go, CCMT – Cambridge Car Memory Task. See Methods for an explanation of the different measures.

### Effect of reliability on correlation between tasks

To test how reliability affects correlations between two different measures in a real-world dataset (such as correlations between brain and behavior, or two different behavioral tasks), we examined different subsamples of our measures, considering two different scenarios: (A) two highly correlated measures, for which both measures had enough data to achieve high reliability, and (B) two measures which were not well correlated with each other, but which both have high reliability (see Supplementary Fig. 2 for correlations between all measures). For each of these scenarios, we subsampled each of the tasks to several different reliability levels (calculated as explained above), and then calculated the correlation between the two tasks. This subsampling procedure was repeated 1000 times to create distributions of possible correlations at a given reliability level (Fig. 3).

**Figure 3.**
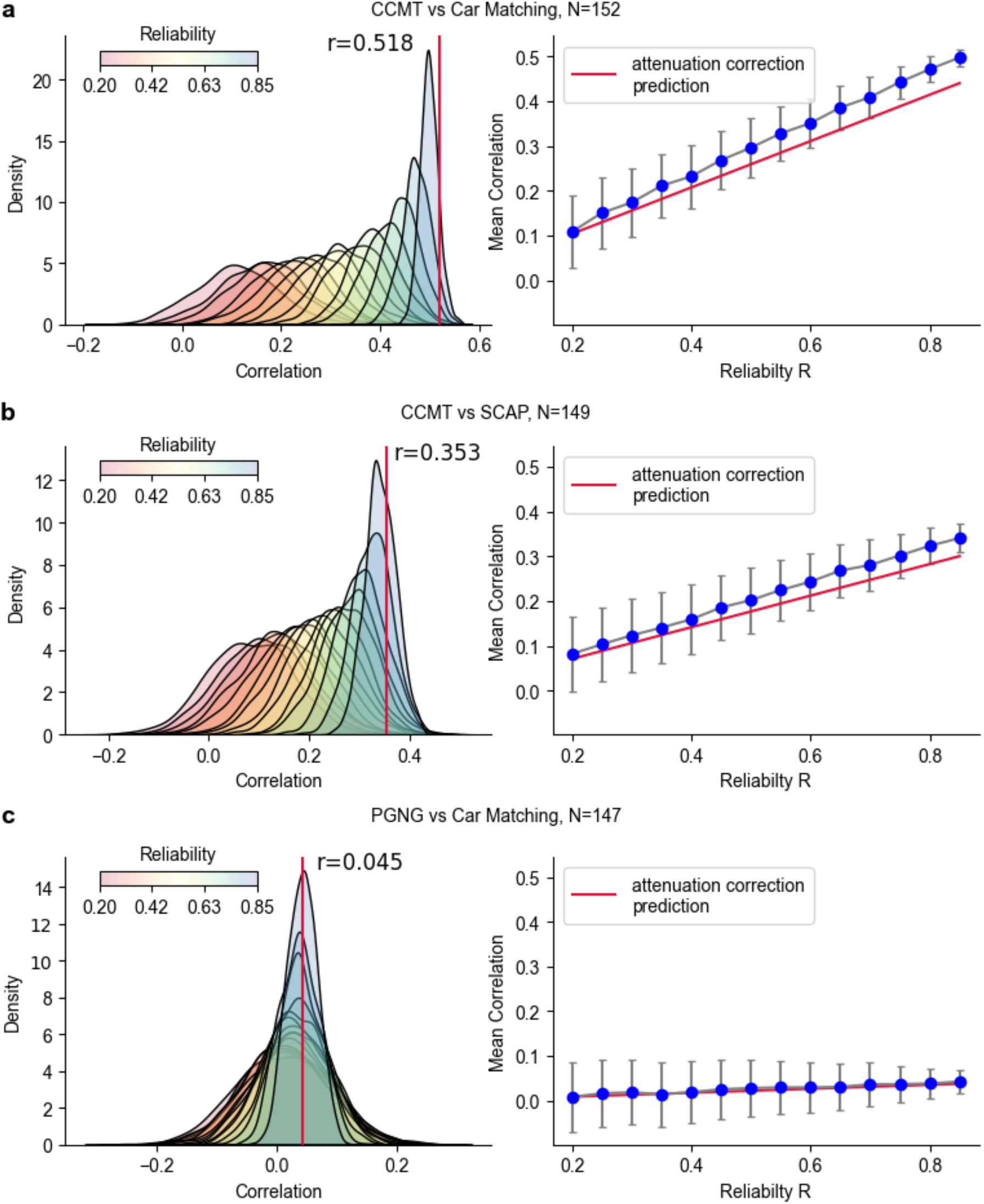
The effect of reliability on correlations between tasks. Left: distributions of correlations between the selected measures colored by resampled reliability of the measures. Red line denotes the correlation between the two measures calculated from the full sample of each. Right: mean correlation over all sub-samples between the two tasks for each reliability level. Error bars denote the standard deviation of correlation values across the different sub-samples. Red line denotes the correlation predicted by the attenuation correction formula. Note that unreliable data generally causes an underestimation of the true correlation values. **a**. Correlation between two highly correlated tasks, the Cambridge Car Memory Test (CCMT) and Car Matching. **b**. Correlation between two tasks with intermediate correlation, CCMT and Spatial Working Memory Capacity (SCAP). **c**. Correlation between two poorly correlated tasks, Parametric Go-No Go Task (PGNG) and Car Matching.

Importantly, low reliability tends to cause an underestimation of the true correlation values. This is in line with the well described attenuation effect first put forward by Spearman^49,63,64^, in which the mean value of the observed correlation is lower than the true underlying correlation, as our data in Figure 3 demonstrates. However, when the true correlation is very low, as in Figure 3b, observed correlations on a single “snapshot” of an experiment, (i.e., one iteration of the split halves analysis), can both underestimate or overestimate the true correlations. Note that overestimation is very unlikely when the true underlying correlation is high.

As reliability of the tasks increases, the distribution of observed correlations between them becomes narrower, as evidenced by the decreased error across the different iterations shown both in the width of the distributions in the left plots, and as error bars in the right plots. The inherent problem with the subsampling method is that with an increased number of trials *L*, we sample a higher percentage of the dataset and thus reduce variance across the distribution of different subsampling iterations. Additional simulations using 10 forms of synthetic data and varying levels of constraining show that the reduced error is not solely driven by the subsampling method as is evidenced by the decreased variance as a function of reliability even for the less constrained data (i.e., when *L* is much smaller than the size of the dataset, see Supplementary Fig. 3).

### Derivation of the fits

Our next question was whether we could mathematically describe the convergence rate of the reliability based on the statistics of the distribution of performance on a given measure. This would allow us to not only calculate reliability for a given dataset and to predict reliability for any given sample size, but would also provide us with a quantification of the convergence rate, an important factor in assessing the suitability of a measure for use in studies of individual differences.

Assuming there is no learning (i.e., samples are independent, and two consecutive trials are independent), and assuming each participant has a true proficiency (i.e., a true ability level for a given task), we would expect participants to each converge to a stable mean reflecting this proficiency. In this case, as we average across more trials, reliability at the individual level will increase, which will drive increasing reliability across individuals (as is indeed shown in Fig. 2 and Supplementary Fig. 1). Under these assumptions, one can derive a formula describing the convergence curve (see SI for the exact derivation):

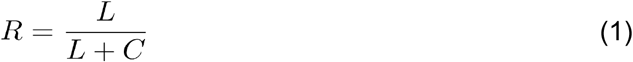

where *R* denotes reliability as defined above, *L* denotes the number of trials used to calculate the reliability and the convergence coefficient *C* is a free parameter, which is fitted to the particular dataset. The maximum sample size is half of the trials, so *L* is always less than or equal to half of the number of trials in the given experiment. Note that this is a reformulation of the classic Spearman-Brown “prophecy” formula^49,50^, which similarly allows the estimation of reliability for larger number of trials based on previously calculated reliability for smaller number of trials^33,51^. This relationship between number of trials and reliability however, not only gives a direct quantification of *C*, but allows a much easier calculation of reliability for any given number of trials.

To further describe the *C* coefficient, and to allow for an even more straightforward calculation of reliability as a function of number of trials in a given task, we sought to describe *C* purely from the statistics of the distribution for that task, without fitting the reliability curve. This is only possible for tasks with binary outcomes on individual trials (e.g., 1/0, correct/incorrect). In these cases, *C* can be expressed using the statistics of proficiencies *f*(*P*):

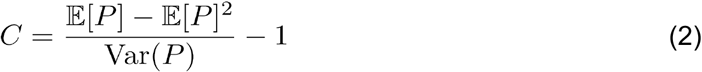

where Var(*P*) denotes the population variance the distribution of proficiencies *P* and 𝔼 [*P*] denotes its mean. In the real world, however, we only have access to the variance of our limited sample and not the true variance of the population. We discuss the impact of limited numbers of participants below. Likewise, for a given participant, we do not have access to their true proficiency, but rather to the proficiency estimated from a limited number of trials. To account for the sampling error which arises from the limited nature of our sampling per participant, we use the law of total variance^65^. We define a random variable *Z* that is an estimate of the participant’s proficiency, leading to the following corrected formula (see SI for derivation):

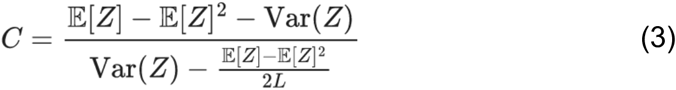

The naive estimation of the *C* coefficient using Eq. 2 leads to a systematic bias which consistently underestimates the *C* coefficient by a large margin (see Supplementary Fig. 4). This bias is entirely corrected even for a small number of trials (*L*>20) using Eq. 3, though a marginal overestimation following the correction remains (Fig. 4, Supplementary Fig. 4).

**Figure 4.**
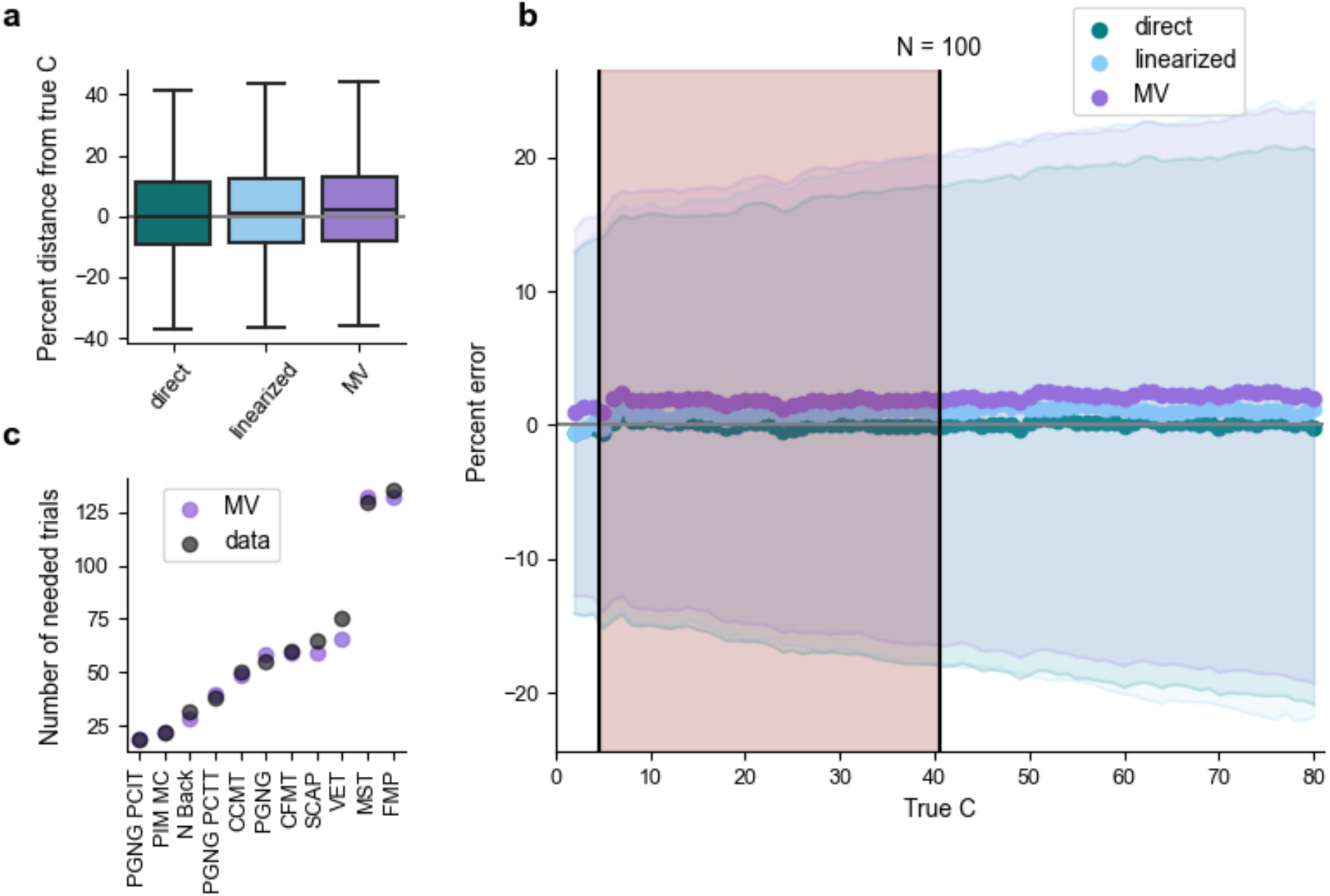
Comparison of three different methods for fitting and estimating the *C* coefficient. **a**. Comparison of fitted *C* value to the true *C* value of simulated beta distributions (absolute difference). Each distribution was sampled 1000 times. Results are shown for *L*=250 trials, *N*=100 participants and for *C* in the range of 2-80. **b**. Error estimation (median and SD of percent error) of the different fitting methods for a range of values of the *C* coefficient (2-80). Data are based on the same simulations used in panel **a**., for *N*=100. The shaded red region denotes *C* coefficient values observed in real-world data (*C* range: 4-41) and the unshaded region shows results from *C* values which we did not observe in our data (42-80). **c**. The predicted number of trials to achieve reliability of 0.8 using the MV fit from Eq. 3 (purple) and an actual number of trials necessary for a reliability threshold of 0.8 as calculated from the data (black) for measures. Included are all measures that in our data reached reliability of at least 0.8.

The *C* coefficient is inversely proportional to the variance in performance on the task across participants – the smaller the variance, the higher the *C*, and the higher the number of trials (*L*) that are necessary to reach a given reliability level (*R*). The *C* coefficient can be fitted from the data using the following three methods (see Supplementary Fig. 4):

1. Directly fitted: fit the convergence curve with hyperbolic function from Eq. (1). Note that this is different from solving the equation for a single given *L* and *R*. To account for noise from a single measurement of *R* (which are reflected in the error bars in Fig. 2a-c) it is advised to fit the full *RL* curve to get *C*.
2. Linearized: fit 1/*R* vs 1/*L* curve, giving a linear fit with an intercept that should be equal to 1 (if trials are independent, i.e., no learning, no fatigue), analogous to Gulliksen^51^:

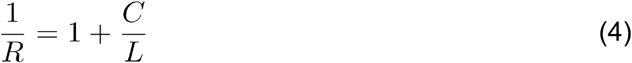
3. Mean/Variance (MV): for binary tasks, fit *C* using the mean and the variance of the underlying *f* distribution using Eq. 3. Note that the first two methods are general and can be used for any dataset, whereas this method is limited to binary tasks, meaning tasks in which the outcome of any given trial is binary and there are only two possible outcomes (0/1, correct/incorrect, blue/red, etc.). Multiple choice tasks with more than two alternatives can be used, as long as the outcome of the trial is binary (e.g., tasks with 4 options, 1 correct and 3 incorrect).

### Validation of the different fits

To compare the different fits and how well they are able to predict reliability, we ran extensive simulations of synthetic data mimicking real behavioral data from our task battery to supplement our analysis of the large real-world dataset we collected. We sampled beta distributions as they are naturally bounded between 0 and 1 and because tasks in our dataset were well approximated by a beta distribution (see Supplementary Fig. 5). We created a dense sampling of *C* values in the range observed in our real-world dataset, and also extended this range further to expand our sampling of the space of potential *C* coefficients. For each distribution, we ran 1000 simulations of an experiment with the same overall *C* (as if participants took the test many times) for several values of *N* (number of participants). We then compared the fitted *C* value from the simulated data to the true *C* value of the distribution from which the data was generated (see SI for the derivation of the *C* coefficient for those distributions). Analyses for both the real-world and the simulated datasets are shown in Figure 4.

As seen from the plots, all three fitting methods (directly fitted, linearized, MV) yield very similar results, and similar (low) error rates (Fig. 4, Supplementary Fig. 4). See Supplementary Figure 6 for an analysis of the intercept of the linearized fit across our real-world datasets, which validates our assumption that there was no significant learning in the tasks included in our dataset. Median percent error was not significantly dependent on the value of *C* (distribution of differences between neighboring points was not significantly different from 0, t_direct, 77_=0.23, p_direct_=0.82; t_lin, 77_=0.82, p_lin_=0.42; t_MV, 77_=0.59, p_MV_=0.56), though the degree of uncertainty in the simulations (SD of the error), did grow as a function of *C* (Fig. 4b). Predictions of the number of trials necessary for achieving 0.8 reliability using the MV fit compared to reliability computed directly from the data show that this error is very minimal even when translated to absolute number of trials (Fig. 4c).

### Dependence of the fits on the number of trials and sample size

To further investigate how a limited number of trials and/or participants might affect the accuracy of our estimate of reliability, we ran 1000 simulations of an experiment with the same range of *C* coefficients as those observed in our real-world data, and varied both the number of participants (*N*) and number of trials (*L*) (Fig. 5). As before for the fixed *N* and *L*, median percent error estimated by the MV fit was largely stable across different *C* coefficients, as shown in Figure 5a. The three values of *C* shown in this plot are representative of values observed in our real-world data: low *C* (left), corresponding to high variance between participants (e.g., Parametric Go-No go Task, percent correct to inhibitory trials), middle range *C*, i.e., middle range variance between participants (middle, e.g., Cambridge Face Memory Test), and high *C*, corresponding to low variance between participants (right, e.g., Mnemonic Similarity Task). As can be seen from all three heatmaps, the median error is negligible and close to zero for *N*>40 and *L*>20. While Figure 5a shows the median error across our simulations, Figure 5b shows the SD of the error calculated across the different simulations. If both the number of trials and the number of participants is very low, error rates are higher, and this is compounded for large values of *C* (reflecting low variance in the data). Because Eq. 3 (which defines the MV fit) is derived for large *N*, it can reach a point of singularity for very small samples (low *L* and *N*), making it impossible to calculate *C*. We remove such rare extreme cases from our simulations (see Methods for details).

**Figure 5.**
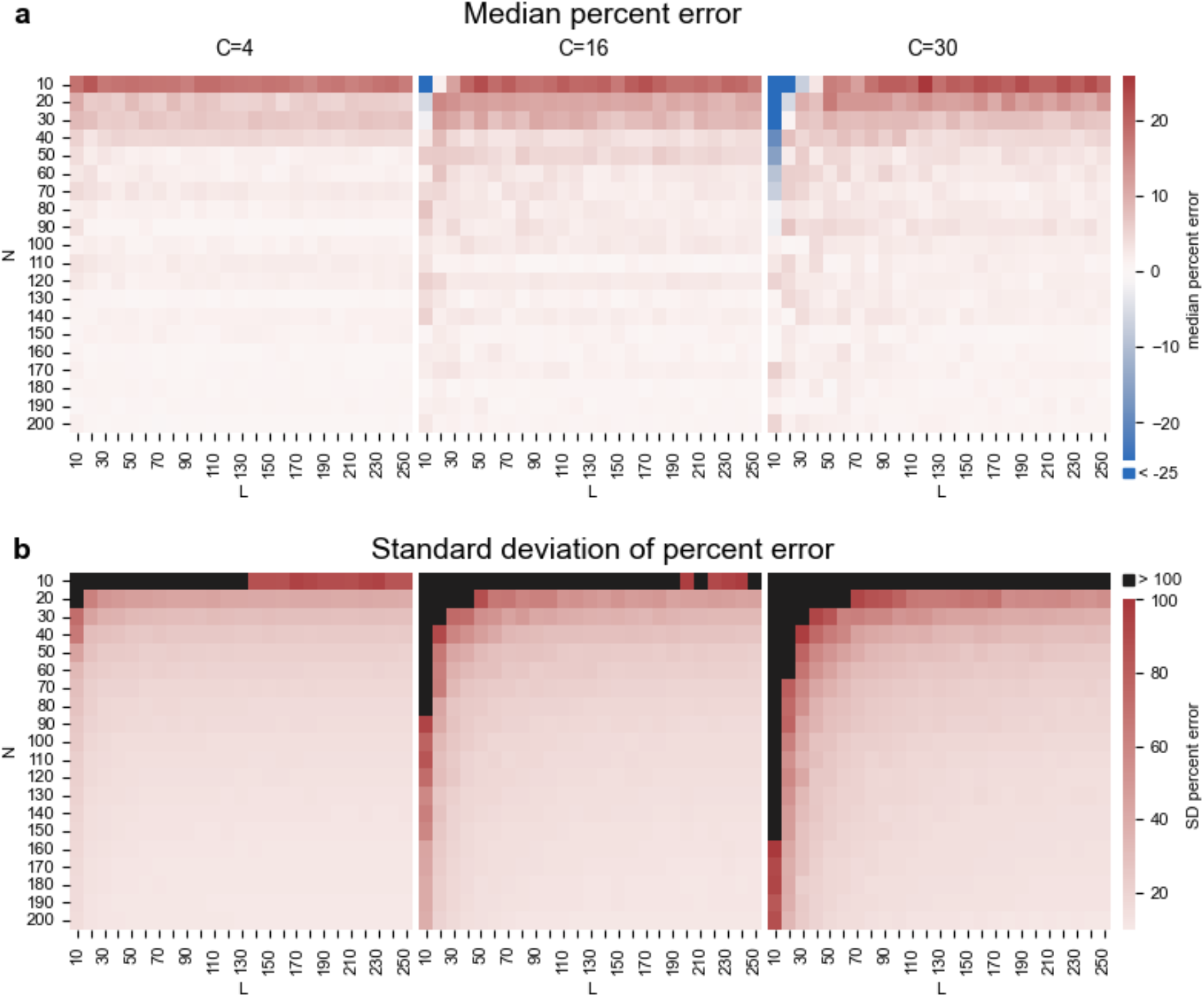
Estimation of error arising from limited sample sizes (number of participants *N* and number of trials *L* per participant) when using the MV formula for selected *C*s matching tasks with low, medium and high variance (PGNG PCIT, CFMT, and MST, respectively) across 1000 simulations. **a**. Median percent error. Range is from -69% to 26% with values below -25% shown in dark blue. **b**. The standard deviation of the error. Range is from 9.8% to 590% with values above 100% shown in black. Note that values over 100% only occur for small *L* and *N* (*L*<20, *N*<30).

### Reliability of real-world tasks

Having established our ability to accurately fit reliability curves to our data, allowing us to predict the reliability for any given sample size, we can now look at the reliability and the projected reliability for larger sample sizes of all the tasks in our battery. Figure 6a shows the number of trials necessary to reach reliability thresholds of 0.8 and 0.9 per task. For analyses which involve correlating different measures, we would recommend 0.8 as a reasonable minimal reliability to aim for, given the effect of reliability on observed correlations (Figure 3). The blue line indicates how many trials there are in the standard administration of each task. As can be seen, while some tasks have enough trials to reach a reliability of 0.8 (blue line above the light brown bar), some are considerably less reliable, and very few have enough trials to reach a reliability of 0.9 (blue line about the dark brown bar).

**Figure 6.**
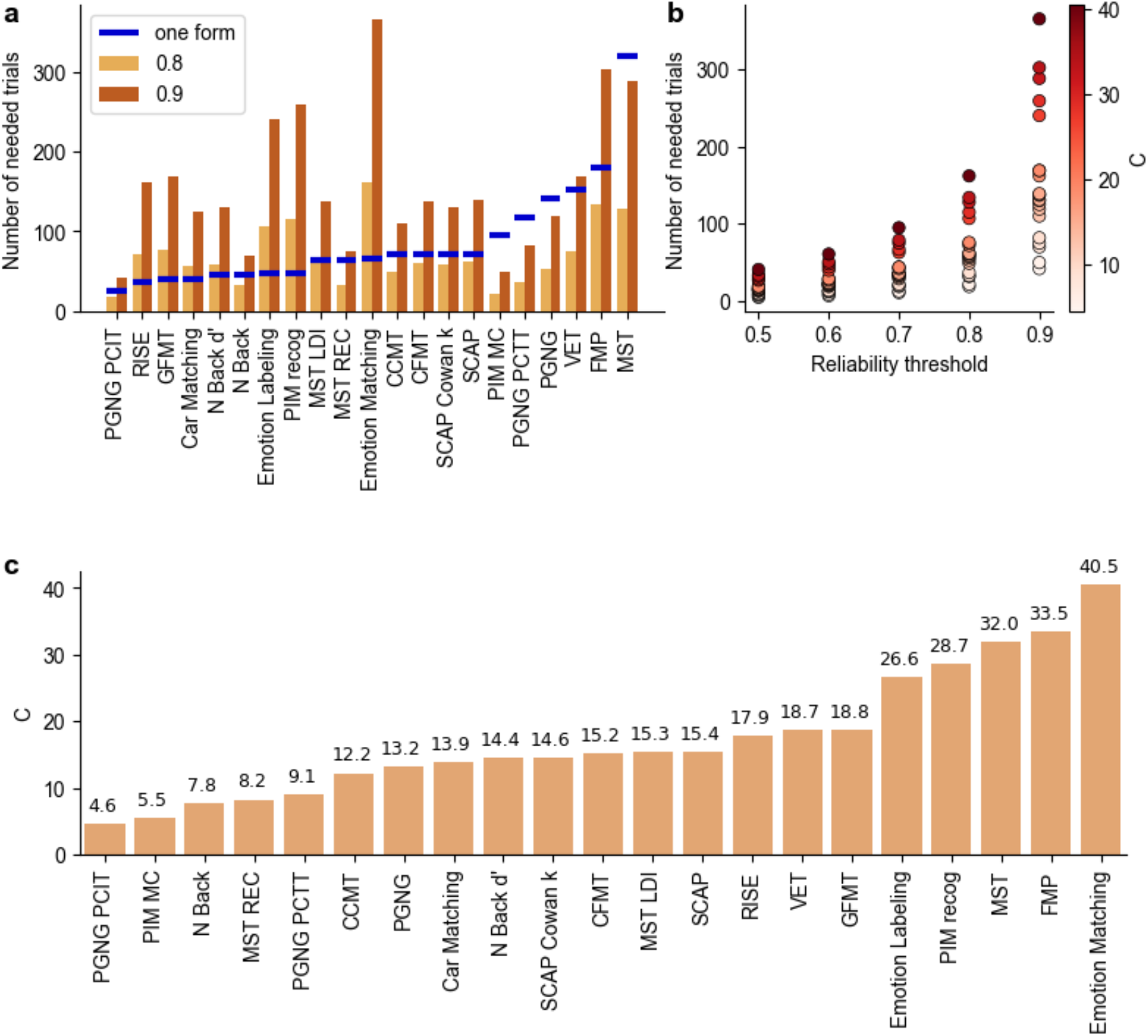
Reliability and its behavior in the real world. **a**. Number of trials needed to achieve given reliability (0.8 in light brown, 0.9 in dark brown) per task. Blue marks denote the number of trials in a single standard form of each task. Tasks where the light brown bar is lower than the blue mark are those where one form shows reliability over 0.8. Predictions were obtained from direct fits of the reliability curves. Note that for some tasks, multiple measures are shown illustrating different aspects of performance. **b**. Dependence of the necessary number of trials to reach different reliability levels in our real-world dataset, colored by the value of the *C* coefficient. Note the non-linear increase in the number of trials needed to reach higher reliabilities, a visual representation of the prediction of Eq. 1. **c**. Value of C coefficient across tasks, calculated from the direct fit. See Methods for an explanation of the different measures.

Figure 6b shows the dependence of the number of trials on both the value of *C* and the desired reliability level. There is a non-linear increase in the number of trials necessary for higher reliability if the variance of the task is low (high *C* values), as predicted by Eq. 1. Figure 6c orders the tasks based on their value of *C*. Note that different measures of the same task can have different values of *C* which would influence their reliability values, even for the same number of trials (e.g., MST REC and MST LDI). For other tasks, different measures might be calculated based on different numbers of trials (e.g. PGNG PCIT and PGNG PCTT). In such cases, it will be difficult to know which measure is more suited for studies of individual differences by simply directly comparing reliability, whereas comparing *C* coefficients will give a clear answer.

### Application of the MV fit for easy study design

Important parameters in study design include the number of trials to be collected and the number of participants needed in order to obtain reliable results at the individual level, especially if the intended use is to investigate individual differences. This is true both for the design of new tasks and for commonly used tasks, which may not have been previously validated for high reliability at the individual level, as was the case for many of the tasks in our battery. In order to provide the community with a tool for building reliable studies, we have developed a simple web application using the MV fit based on our results above. To use this app, a researcher could run an initial pilot study using a small number of trials, which will be used to estimate the mean and sample variance (across participants) of the population. These numbers, along with the number of participants on which they were calculated, can then be entered into the app, which will output an estimate of the necessary number of trials to reach any given level of reliability.

To account for inaccuracies due to low sampling of the task’s distribution resulting from small sample sizes, we implemented the error estimated from our extensive simulations (Fig. 4b, Fig. 5) in the app, as confidence intervals.

The tool is available at https://jankawis.github.io/reliability-web-app/, and its basic functionality is depicted in Figure 7 (see SI methods for use instructions).

**Figure 7.**
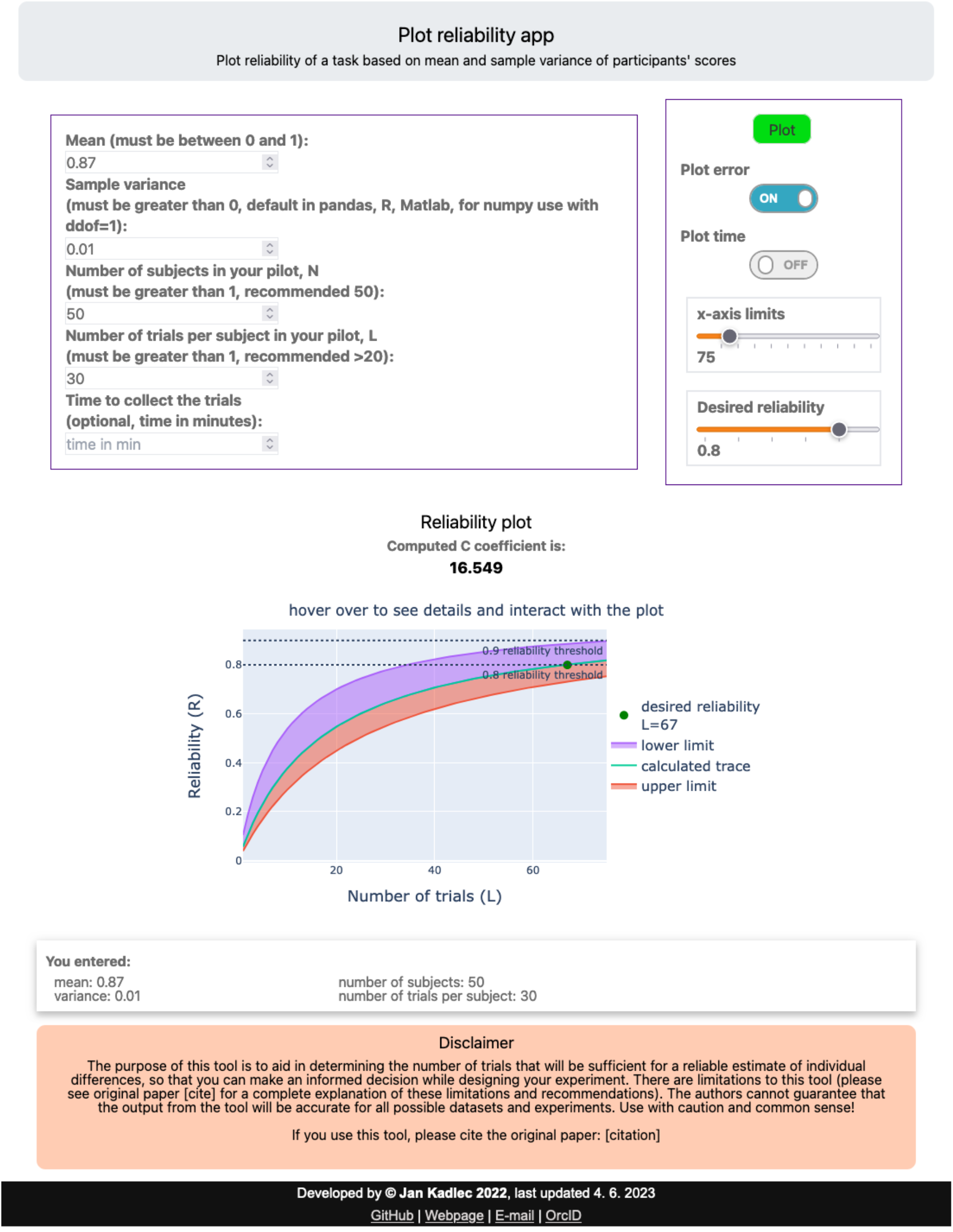
Web application for reliability estimates.

## Discussion

As the focus in cognitive neuroscience research in recent years has partly shifted towards precision medicine and to understanding individual differences, it has become crucial to better validate behavioral measures to ensure high reliability at the level of the individual. A common approach adopted in many studies is to use individual differences in behavioral measures to search for the neural variance which explains them, i.e., brain-behavior correlations. In many of these studies, behavioral measures are collected separately from the neural data, often at a different time point, for many good practical and technical reasons. Since the behavior-neural comparison in such cases assumes that the behavior has not changed in the interval, the trait-like stability of the behavioral scores at the individual level is likewise critical for such analyses. Yet the very existence of such stable, trait-like individual performance across days has mostly been assumed rather than directly demonstrated in many commonly used tasks. Moreover, many cognitive measures were designed to detect group effects, where the reliability of scores per individual is not as important^47^. Averaging over many participants achieves the same effect of convergence to a stable group mean, even if individual reliability is low. Here we set out to directly test the assumption that relative individual performance on many different commonly used tasks will converge to a stable average given sufficient data, and to ask the more complex question of exactly how much data is sufficient and whether it is possible to estimate this value before full data collection, rather than after.

As was previously reported in the literature^40–46^, and can also be seen in our data (Figs. 2 and 6), many tasks have only modest reliability at the individual level. The importance of high reliability for detecting true correlations has also been previously discussed, and the effects of low reliability have previously been shown in simulated data^31–33,36,51,66–70^. We demonstrate these effects on real-world data in Figure 3 which shows how reliability affects the observed correlation between two different measures, grossly underestimating high correlations, and either underestimating or inflating very weak correlations. In this manuscript, we examine correlations between behavioral measures, but the same effect should be seen for low reliability on the correlation of any two measures, including brain-behavior correlations (see also Nikoladis et al.^32^ for a more in-depth discussion of the effect of behavioral reliability on brain-behavior correlations, which is mathematically similar to comparing two behavioral tasks). Our data are therefore reassuring in the context of the debate regarding inflated correlations in brain-behavior analyses, if moderately high correlations are observed (for instance Figure 3b shows that a correlation of 0.35 is very unlikely to be overestimated).

There are many different definitions of reliability, which are used for different purposes. Reliability, as we define it here, is an expansion of the traditional test-retest reliability which involves administering the same task twice to the same group of participants, and then calculating the Pearson’s correlation between the two repetitions. This reliability, which is focused on the internal consistency of the scores across participants, rather than the absolute scores themselves, is relevant for our aim of improving the reliability of individual differences measures. However, in cases where the aim is to validate the consistency of the absolute score on each of the items on the test in order to provide population norms and statistics, subsampling of different trials within the task as we do here is not helpful, and a different calculation of reliability is used^68,71–73^. For a detailed discussion on these two divergent and often confusing definitions of reliability, see Snijder et al.^37^

How can we determine which tasks are best suited for studies of individual differences? This has become an even more pressing issue with the advent of online testing, which has promoted the development of many new task paradigms. A tool which could both inform study design by quickly and easily predicting from small amounts of pilot data the necessary number of trials to achieve high reliability, as well as provide a straightforward evaluation and comparison between tasks of their efficiency for individual differences studies, would therefore be of great value to the community.

To address these challenges, we mathematically derived a simple equation which is a reformulation of the Spearman-Brown prophecy. Our equation introduces the convergence parameter *C*, that describes the rate of convergence of the reliability curve as a function of the number of trials for a given measure. *C* is inversely proportional to the variance in performance across participants – i.e., the lower the variance across participants, meaning the narrower the distribution of performance, the larger the value of *C*, and the more trials will be necessary in order to achieve reliable separation of individuals. Intuitively, tasks with very low variance in performance across the population, for instance, due to floor or ceiling effects, will need a large number of trials in order to rank participants reliably and maintain internal consistency. This relationship between *C* and the number of trials needed for increased reliability is non-linear, as is shown in Figure 6b.

Our analysis is unique in that we not only validate our calculation of the *C* coefficient on simulated data, but also on a broad set of real-world behavioral datasets across many different tasks and cognitive domains. Crucially, all these data across all the different tasks were collected on the same participants, allowing us to directly compare the tasks (Table 1). Simulations are often the gold standard as they are not limited by expensive and time-consuming data collection, but they require making assumptions on the data which may not hold true for different tasks. Figure 4 shows that applying both the direct/linearized fit formulas as well as the MV fit very accurately predicted the reliability for different sample sizes across both our real-world dataset as well as across large simulated datasets. Figures 4c, 6 and Supplementary Figure 1 show that different tasks had very different convergence rates, even when measured within the same individuals.

The remaining question is how much data are needed in order to get a reasonable estimate of reliability and of the *C* coefficient, for instance in the case of a pilot study. The accuracy of the estimate depends on both the quality of the data and the sample size. When collecting data online, it is recommended to first thoroughly clean the data, removing non-compliant participants and identifying participants who were inattentive to the tasks^74–76^. This is, naturally, also true for data collected in the lab. As for the effect of sample size on the potential error in the estimation of *C*, we ran extensive simulations to estimate the dependence of the error on both the number of participants (*N*) and the number of trials (*L*) (Fig. 4b, 5). A surprising result from this analysis is that beyond a relatively small number of participants (*N=*∼40-50) and a small number of trials (*L*>=20), the reliability estimates are very robust, and the error is not greatly reduced by increasing sample size. For a given number of trials, this is achieved in our formula through the implementation of the law of total variance, which corrects for the systematic underestimation of *C* which exists otherwise (Supplementary Fig. 4). Our results suggest that collecting data from 40-50 participants will provide a reasonable sampling of the statistics of the population for these tasks, and also serves as a general guide to selecting the minimum number of participants for an individual differences study.

With regards to brain-behavior association studies, one strong recommendation that emerges from our study is to focus on highly reliable tasks, and consider the tradeoff between data per participant and number of participants. By collecting more data per participant, using highly reliable tasks, experiments could preclude the need for thousands of participants. The other recommendation would be to focus on behavioral tasks and neural measures that do not have very low correlations between them (r<0.1), unlike many of the brain-behavior correlations discussed by Marek et al.^23^ While not guaranteed, higher correlations could be accomplished by choosing tasks that are more domain-specific (and thus likely to better predict the connectivity strength between certain nodes), by using multivariate instead of univariate neuroimaging measures, or by functionally defining relevant regions individually instead of using generic anatomical parcellations.

Although our real-world dataset collected online represents a more heterogeneous population than the typical study conducted in-person in a lab (and generally overwhelmingly composed of college students), it is still likely less heterogeneous than the full general population. It is possible that more participants will be required to reach target reliability thresholds for the tasks we report here when studies include more diverse populations, such as clinical populations.

Our reformulation of the Spearman-Brown, along with the derivation of *C* as a function of the mean and variance of the distribution, provides a simpler way than was previously available of calculating the reliability of any given dataset, and predicting how many trials will be needed to reach any given level of reliability. Perhaps more importantly, this reformulation suggests another way of conceptualizing reliability not only as an outcome to strive towards, but also as a core, measurable property describing the suitability of the task for individual difference studies. The convergence parameter *C* provides us with a “score” of how well a task can reliably separate individuals, above and beyond the reliability of the standard administration of that task which could vary with number of trials included in the form (Figs. 4 and 6). It is a direct assessment of the task itself as a measure of individual differences, not of how well the standard administration does in collecting sufficient data in terms of having enough trials. This can help evaluate the suitability of that particular task for individual differences studies in relation to other similar tasks. For instance, the standard administration of the Cambridge Face Memory Test (CFMT) is more reliable than the Glasgow Face Memory Test (GFMT), and also more suitable for studying individual differences (lower *C*). Emotion discrimination tasks (emotion labeling and emotion matching) have very low reliability for a standard administration. These tasks also have high values of *C*, perhaps pointing to a lack of objective answers for these tasks. We provide the range of *C*s calculated from our dataset as a starting point (Fig. 6c), to help relate new tasks to an existing, though partial, scale. Note, however, that tasks can differ greatly not only in how many trials are necessary to reliably separate individuals but also in how long it takes to collect said number of trials, time being a key practical parameter in real-world studies. As the dependence of the number of trials on *C* is not linear, there is no trivial comparison between two tasks with different *C* parameters in terms of the time taken to collect the data. Experimenters are therefore advised to factor in the time necessary to collect enough data for their desired reliability level.

Our results show that reliability of standard tests should not be implicitly assumed but rather experimenters need to explicitly ensure that the test instrument, together with experiment parameters including number of subjects and number of trials, collectively yield a study with acceptable reliability. The analytical tools we presented in this study can be adopted as a general tool for study design, and can be used both to project the necessary number of trials/participants for any desired reliability level, based on only a pilot study as well as to characterize reliability of existing data sets (e.g., before subjecting them to brain-behavior association analyses). In order to maximize the accessibility of this tool, we created a simple web application that allows researchers to obtain the reliability curve and the *C* coefficient simply by entering the mean, sample variance, number of participants and trials in their study. The application also has the option to calculate the time it would take to collect enough trials for any given reliability level by including the amount of time taken to collect a particular sample (Fig. 7). The web app implements confidence margins, based on the mean and standard deviation of the error observed across our widespread simulations. This addresses the potential for a small error in the calculated *C* leading to a large error in projected number of trials, as discussed by Charter^61^. Our hope is that by reducing demands on the computational background of end-users to an absolute minimum, the web app will remove any barrier that would otherwise hinder the applicability of this finding and allow researchers across different fields to easily use this tool in their research to optimize study design.

Note that the only assumption we have made in deriving these fits is that there is no learning, or alternatively, fatigue over time as participants perform each task. To minimize fatigue, we split our tasks across several shorter testing days, providing participants with a financial bonus for completing all tasks across the different days in order to minimize attrition. To ensure that there is no learning, we used alternate forms whenever more than one form was collected (except for the CCMT, which shows no learning even with the same form, as is shown by the intercept for the linear fit, Supplementary Fig. 6). For most tasks, the use of alternate forms is important to counteract learning. It follows from the equation of the linearized fit (Eq. 4) that if trials are independent (an assumption in our derivation which would translate to no learning or fatigue), the intercept of the fit must be one, which is indeed the case for all or most of our measures (see Supplementary Fig. 6). Deviations from an intercept of 1 on the linearized fit suggest a violation of the assumptions (no learning / fatigue). This could provide a way of testing the presence or absence of learning or fatigue in a particular dataset, as well as opening up potential directions for future explorations of learning by developing further models to understand the interaction between learning and the intercept.

In summary, we validated our reformulation of the Spearman-Brown prophecy in a range of real-world cognitive tasks across many different cognitive domains, showing that the reliability of these tasks does eventually converge. We conceptualize this convergence coefficient *C* as a fundamental trait of a given task, which can be used to evaluate its suitability for testing individual differences. Importantly, we provide the community with a simple online tool that computes the *C* coefficient, projects the reliability curve using only descriptive statistics of the experiment (mean, variance, number of participants and trials), and can be easily used to aid in designing new studies.

## Methods

### Real-world behavioral tasks

Since both the tool and the derived formula are as general as possible, and in order to make this tool widely applicable, we tested it on known tasks across many domains to cover a wide spread of cognitive measures. Included tasks are detailed in Table 1 (including average time to completion and number of participants with one and more forms), and measures are summarized below. For some of the tasks, we examine multiple measures which reflect different aspects of performance (e.g., inhibitory control vs. sustained attention). Additionally, we provide detailed descriptions of the tasks and measures with examples and references at https://jankawis.github.io/battery_of_tasks_WIS_UCLA/intro.html.

#### Working memory

- Spatial Working Memory Capacity (SCAP)^77^: Cowan’s K^78^ (maximum across loads), and overall accuracy
- Parametric Go/No Go Task (PGNG)^40,79^: percent correct to target (PCTT), percent correct to inhibitory trials (PCIT), calculated across levels, and overall accuracy
- Fractal N-Back^80^: d’ for 2-back, and overall accuracy calculated across levels

#### Object memory

- Cambridge Car Memory Test (CCMT)^81^: overall accuracy
- Vanderbilt Expertise Test (VET; collapsed across birds, leaves and planes subtests)^82^: overall accuracy
- Mnemonic Similarity Task (MST)^83^: recognition accuracy score (REC), lure discrimination index (LDI) and overall accuracy. Note that here we used the traditional version of the MST; a new version (oMST^84^), which we did not test, is now available.

#### Object (car) perception

- Car Matching Test^85^: overall accuracy

#### Object/associative memory

- Relational and Item-Specific Encoding (RISE)^86^: overall accuracy on relational blocks. Note that we didn’t include the repetition in the analyses due to its poor reliability (*R*=0.26).

#### Face memory

- Personal Identity Memory Task (PIM): overall accuracy on multiple choice attribute recollection test (PIM MC); face recognition accuracy scaled by confidence (PIM recog)
- Face Memory/Perception Task (FMP): overall accuracy

#### Face perception and memory

- Glasgow Face Matching Test (GFMT)^42^: overall accuracy
- Cambridge Face Memory Test (CFMT): overall accuracy; data included from original^2^, Australian^87^ and Female^88^ test forms

#### Social cognition

- Emotion Labeling and Emotion Matching Tasks^89^: overall accuracy

In order to acquire a sufficient number of trials per participant to estimate the true reliability of each task and correlations between tasks, we either administered the task multiple times (SCAP, CCMT), administered extended task versions that included additional trials with novel stimuli (GFMT, Emotion Matching and Emotion Labeling) or administered additional versions of the task with novel stimuli (MST, PGNG, RISE, PIM, FMP, CFMT, VET).

### Data collection

All data were collected online and participants were recruited using the online platform Prolific (www.prolific.co). 298 participants started the first day of the experiment. Out of them, a total number of *N*=257 (131 females, 120 males, 6 not stated, mean age: 29.8 ± 7.7) finished all tasks of the first experimental day and were included in our analyses. Of those, 244 completed the full 3-day battery. Participants were paid at a rate of $9.50/h, plus an additional $9 if they finished all the tasks within the week. Eight months after successfully completing the full battery (see below for exclusion criteria per task), we invited participants back to increase the power of our analysis and investigate reliability. At this stage we introduced two additional tasks to the battery – the Vanderbilt Expertise Task (birds, leaves and planes subscales) and a visual N-Back task with fractal stimuli, which included 0-back, 1-back and 2-back conditions. A total of *N*=183 returned and did some portion of the tasks (see Table 1 for counts of participants per task). 89 participants successfully completed all tasks with at least one repetition. Of these, 41 participants successfully completed all the repetitions.

### Inclusion/Exclusion criteria

We chose to exclude participants on a per-task basis, based on a combination of their accuracy, reaction time (RT) and individual trial responses. We assessed the following criteria and excluded participants from a task if two or more of the following criteria were met:

- Average RT was 2 standard deviations (SD) faster than the group mean.
- Standard deviation of RT was less than 2 SD below the standard deviation of the group.
- Average sequence length of a single response was 2 SD greater than the group mean sequence length.

If a participant’s accuracy was greater than 0.5 SD below the mean, they were included, regardless of their RT and individual trial responses. The only exception to this was for the MST, where if the standard deviation of the RT was less than 2 SD below the SD of the group, they were excluded regardless of performance, as this pattern of responses suggested that the task was being performed by a script/bot rather than a human.

After excluding participants based on accuracy, RT and individual trial responses, we additionally excluded participants based on the following criteria, which would indicate that they were not paying attention during the tasks:

- CFMT: Four or more incorrect trials in Stage 1 (i.e., Stage 1 score less than 83%). For participants with three incorrect trials in Stage 1, data were excluded if performance on the other stages indicated a lack of attention rather than a valid measure of poor performance.
- FMP: Accuracy below chance (50%) in face-matching trials.
- PGNG: Accuracy less than 3 SD below the mean for the two target identification stages. If accuracy was less than 2 SD below the mean, performance on the rest of the task was evaluated to determine whether low accuracy was because of a genuine lower performance or lack of attention.
- N-Back: Accuracy less than 3 SD below the mean for the 1-back blocks. If accuracy was less than 2 SD below the mean, performance on the rest of the task was evaluated to determine whether low accuracy was because of a genuine lower performance or lack of attention.
- VET tasks: Incorrect or missing responses on 2 out of 3 catch trials.

## Data analysis

### Reliability calculations

Reliability was calculated using the following algorithm (Fig. 1): Define batch sizes (*L*) of trials from *L*=1 to half the total number of trials. Then, select two sets of non-overlapping *L* trials for each participant, compute the outcome measure of interest, and calculate Pearson correlation *R* between those two vectors of length *N* (number of participants). Repeat this sampling 1000 times per each *L* and use the mean of these *R* values in the *RL* plots.

The procedure is described in the Results in the section on measuring reliability and depicted in Figure 1. We tested different numbers of samples ranging from 100 - 10,000 and chose 1,000 as that was sufficient for stable results without being too computationally costly.

In order to determine whether random data would show spurious but reliable correlations given a large enough number of trials, we simulated a set of random data. We randomly generated zeros and ones for *N*=100 participants with *L*=250 trials, and calculated reliability as we did for the real data. As different simulations gave reliability values that ranged between -0.1 to 0.1, we tested whether there was any consistent reliability in the random data by repeating the simulation 100 times and averaging the reliability curves. We then used a student’s t-test to test whether this averaged reliability curve was significantly different from zero, and it was not (two-tailed t-test, t_99_=1.17, p=0.25).

To calculate reliabilities for our battery of tasks (Fig. 2d), we used two forms of each task to get a reliability that corresponds to one commonly used form of each task. For tasks that indexed memory (i.e., where repeating the task with the same exact stimulus set would potentially create learning-related improvements due to additional exposures), the additional versions of the task used the same task structure but novel stimulus sets. Specifically, for the CFMT, we used the original form^2^, and as a second form, we used the Australian version^87^. For VET tasks, three subtests (planes, birds, leaves) were administered and trials were concatenated to create one full form. For tasks where repeating the same task would not create any learning-related improvement, such as the working memory tasks, the same task was administered twice.

### Effect of reliability on correlations between tasks

For the analysis in Figure 3, estimating the effect of reliability on the correlation between two tasks, we used the following approach. From our battery, we selected two pairs of measures. The first pair, CCMT and Car Matching, have high reliability within each measure for all trials and also high correlation between the two measures (r = 0.518). The second pair, Car Matching accuracy and PGNG overall accuracy, have high within-participant reliability of at least 0.85 for each measure separately (when using all the available trials), but a low correlation between the two measures (r = 0.045). For each task, we calculated how many trials are needed to achieve different reliability values – from 0.1 to 0.85. In order to be able to utilize the full dataset for estimating the correlations, the number of trials per reliability level (per task) was calculated using the directly fitted fit method (see Results - derivation of the fit).

Having determined the necessary number of trials for each reliability level, we then subsampled the corresponding number of trials from each of the tasks and calculated the correlation between each of the two task pairs. We repeated this 1000 times to create a distribution of correlations between those two tasks for a given reliability level, using the appropriate number of samples. The right panels of Figure 3a and 3b display the mean and standard deviation of the correlation distribution for a given reliability (R).

### Error estimation

To estimate error (Fig. 4 a,b) for validating our method, we created synthetic data sampling *C* coefficients from 2 to 80 by 1 (79 values). We used a beta distribution that is defined using two parameters, *α* and *β*. We verified that all measures in our dataset can be well approximated and fitted using the beta distribution (Supplementary Fig. 5). It can be derived (see SI) that *C* = *α* + *β* and we used *α* = *β* when generating beta distributions for simplicity. For a given value of *C*, the value of *α* would uniquely specify *β*, with a different *α / β* ratio. We tested that the ratio of these coefficients does not make a substantial difference when comparing the error of Eq. 3 (see Supplementary Fig. 7).

For each value of *C*, we ran 1000 simulations (realizations of each given combination of mean and standard deviation). When calculating reliability to obtain *RL* curves for directly fitted and linearized fit, we sampled *L* only 500 times to reduce computational cost. Each dataset contained 250 trials per participant for 20 different values of *N* (number of participants) ranging from 10 to 200 by 10 to create scaling of error with *N* (Supplementary Fig. 8). The ground truth for the *C* coefficient was established for each *C* separately by generating a distribution with large sample size of *N* = 10^7^ and using MV fit to get the *C* corresponding to this distribution. For all three fits, we computed the difference between the fitted *C* and this true *C*. We then divide this distance by the true *C* to get error in percent and report the median and standard deviation of this percent error (Fig. 4 a,b).

To prevent artefactual error arising when the denominator of Eq. 3 approaches zero (happening for low values of *N* and *L*), we disregarded simulations where the denominator was less than 10^−3^. This threshold was determined by finding inflection points and maximas of second derivatives of the MV fit function. That yielded a range of thresholds from 3*·*10^−3^ to 10^−3^ and the less stringent threshold was chosen as a compromise between the possible error and the probability of such an event (computational cost).

The same approach with the same *C* values and policies was used to generate the error matrices in Figure 5. The only difference is that each of these *N* and *C* combinations was created for 25 different *L* values ranging from 10 to 250 by 10 and only the MV fit was used. We compute the percent error as the median (Fig. 5a) and standard deviation (Fig. 5b) across the 1000 simulations of the distance of the fitted MV value from the true *C* value scaled by the true *C* value. This yields a *N*x*L* matrix of percent error for each *C*.

### Software and used packages

All the analyses were performed using Python 3^90^ and the following packages numpy^91^, pandas^92^, matplotlib^93^, seaborn^94^, lmfit^95^, scipy^96^. Final figures were created using the CanD^97^ package. The web app is designed using pyscript^98^, HTML, CSS and the same python packages as mentioned above.

All the tasks were coded using lab.js^99^ (www.lab.js.org) and run on our servers.

## Code and data availability

Code and data will be made accessible on GitHub at the time of publication.

## Acknowledgements

We thank Sasha Devore for insightful comments during the writing process. We would also like to thank all participants that took part in this study. This work was generally supported by ISF grant 829/22, and the Zuckerman STEM leadership program. M.R. is the incumbent of the Roel C. Buck Career Development Chair.

## Supplementary figures

**Supplementary figure 1.**
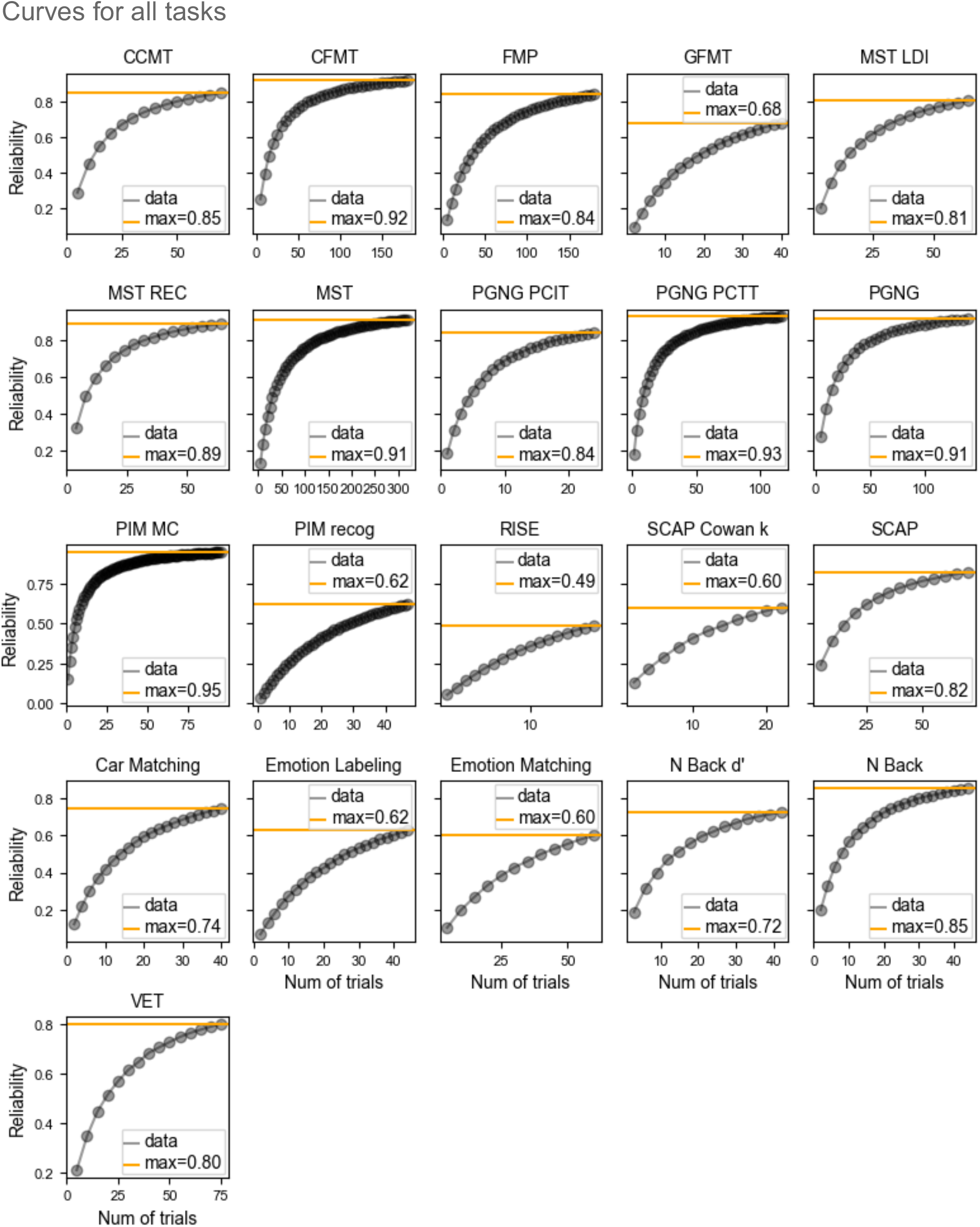
Reliability curves for all behavioral measures (21) in our battery.

**Supplementary Figure 2.**
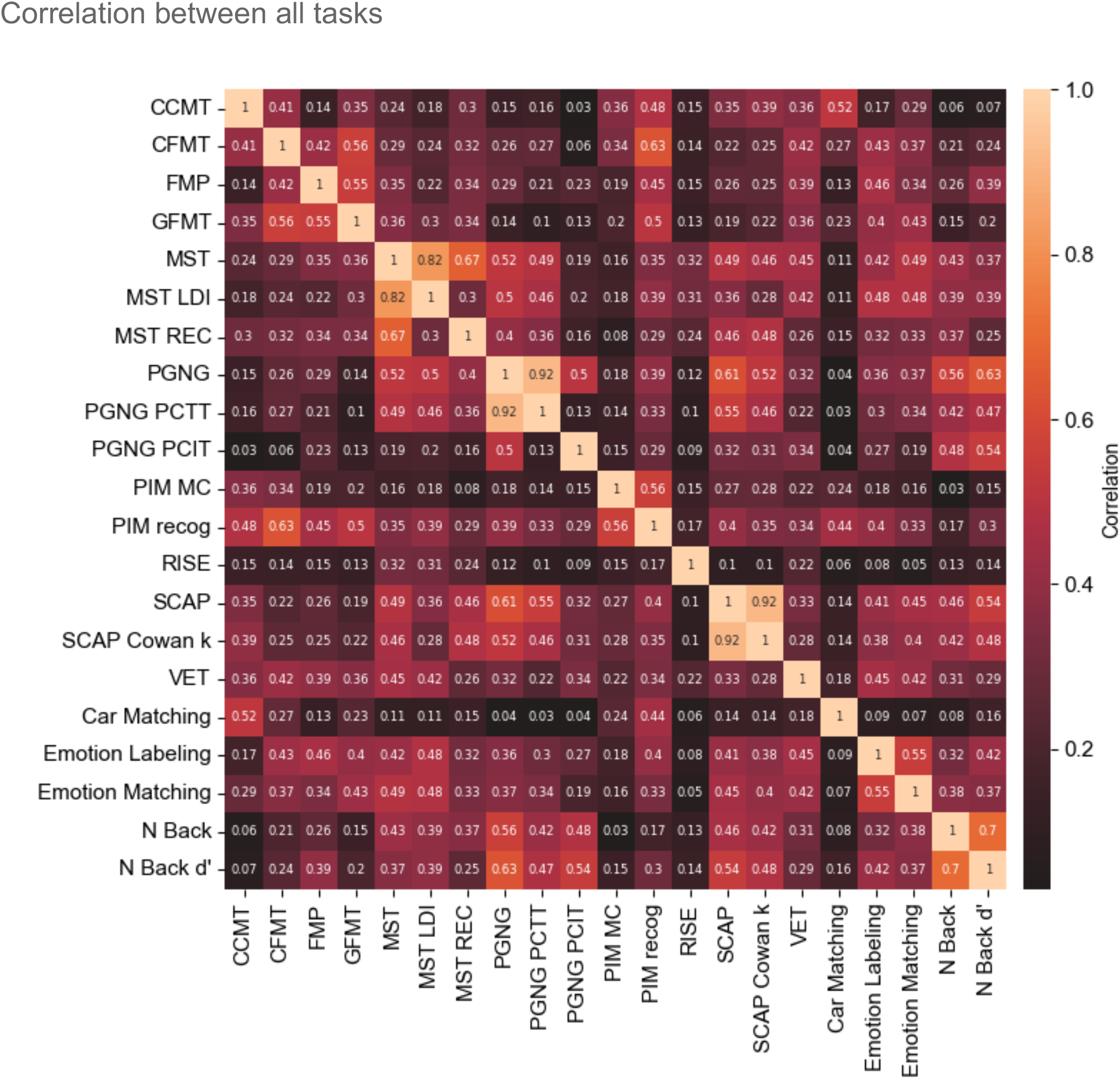
Correlation between measures across all tasks (14) and measures (21).

**Supplementary Figure 3.**
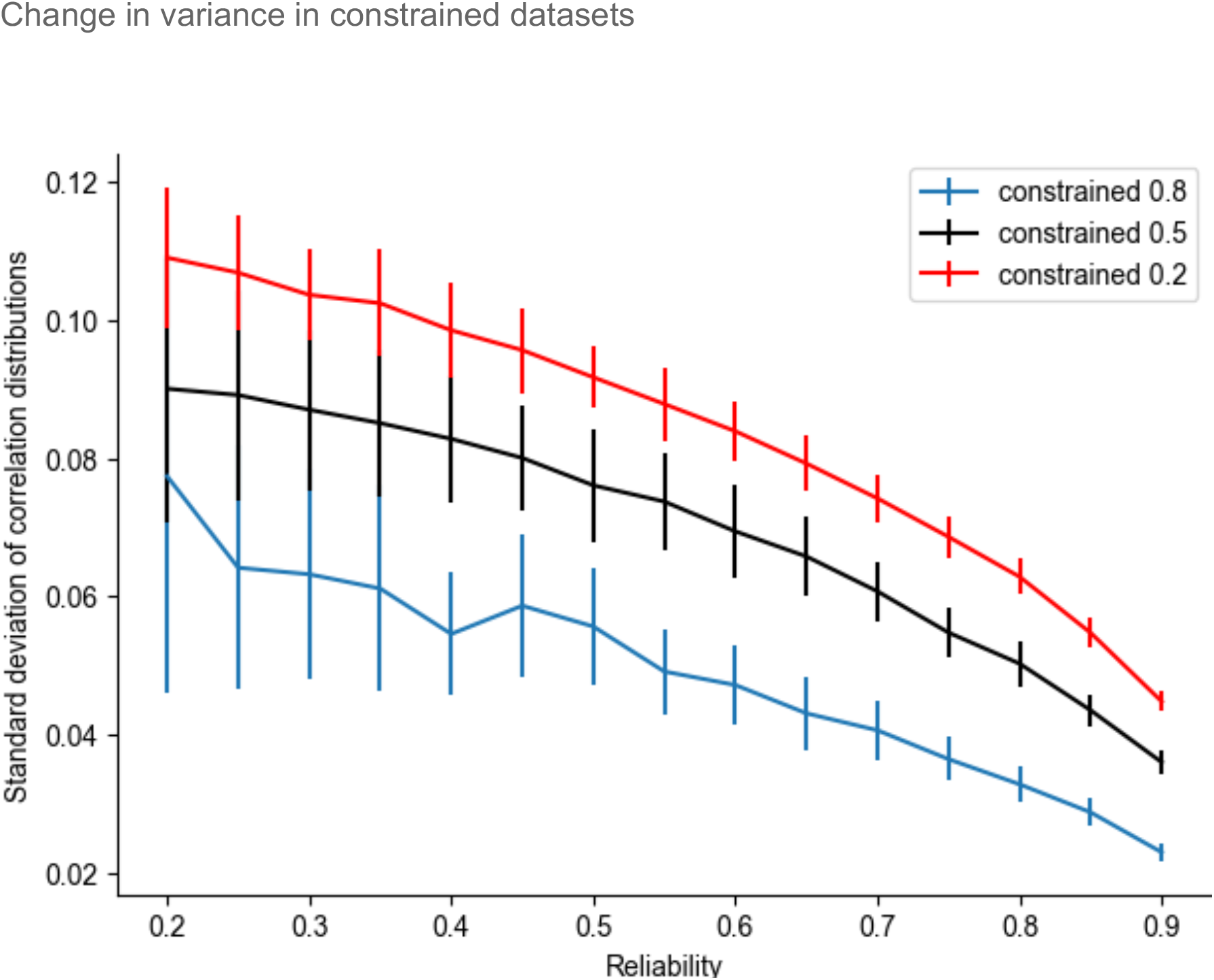
Mean standard deviation of correlation distributions for reliability level across 100 simulations of 10 forms of synthetic datasets based on statistics from our real-world tasks for *N*=75. Colors show different levels of constraints of the dataset (i.e., what percentage of the dataset the *L* trials used to calculate reliability consisted of) when calculating reliability to ensure the reduction of standard deviation with increased reliability was not an artifact of decreased variance in the subsampling. Standard deviation decreases with increasing reliability (regardless of the size of the dataset that trials were sampled from), suggesting that the correlation between the tasks is truly better estimated and not the result of an artificial reduction in variance.

**Supplementary Figure 4.**
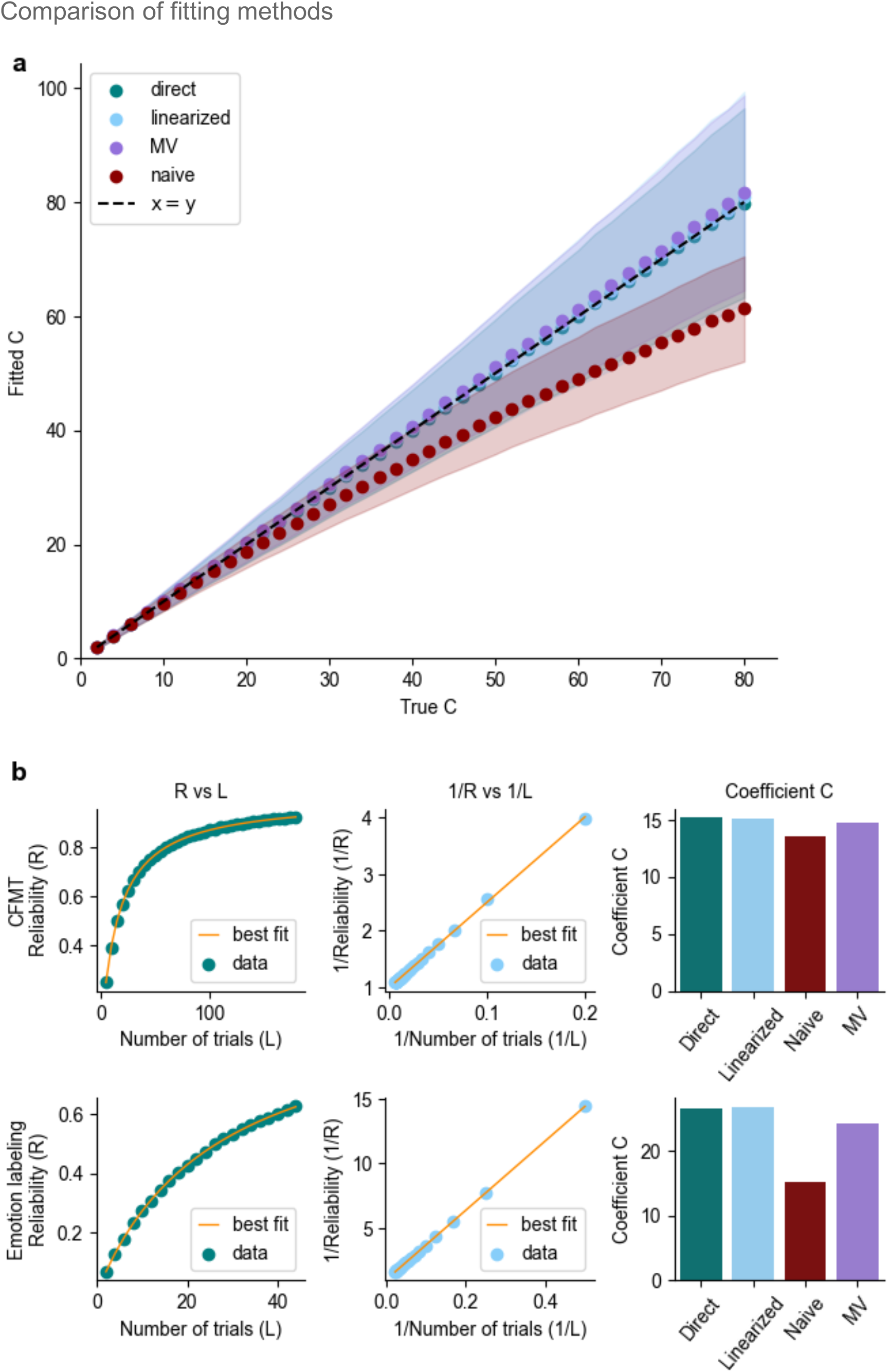
**a**. True *C* versus fitted *C* using four different methods derived and discussed in the paper for *N*=100 and *L*=250. Naive method (red) used to fit the data using Eq. 2 consistently underestimates the *C* coefficient. **b**. Comparison of C estimations using different methods for CFMT (top) and Emotion labeling (bottom) tasks. First column shows the direct fit (Eq. 1), middle column depicts linearized fit (Eq. 4), and the right column shows *C* values obtained for each of the four fits (Eq. 1-4).

**Supplementary Figure 5.**
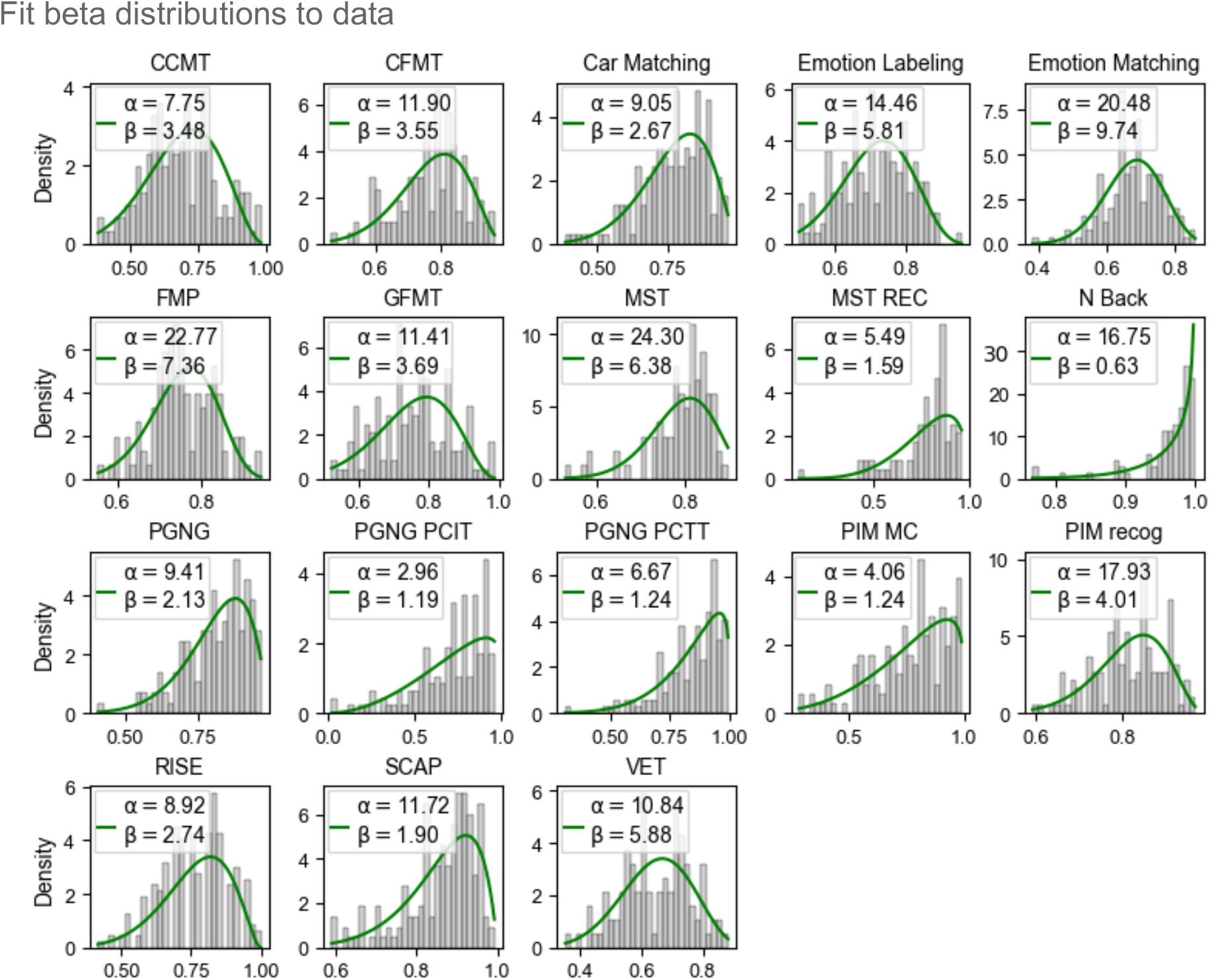
Fit of beta distributions to all relevant measures (18). Barplot shows the distributions of scores and the green line shows the best fit.

**Supplementary Figure 6.**
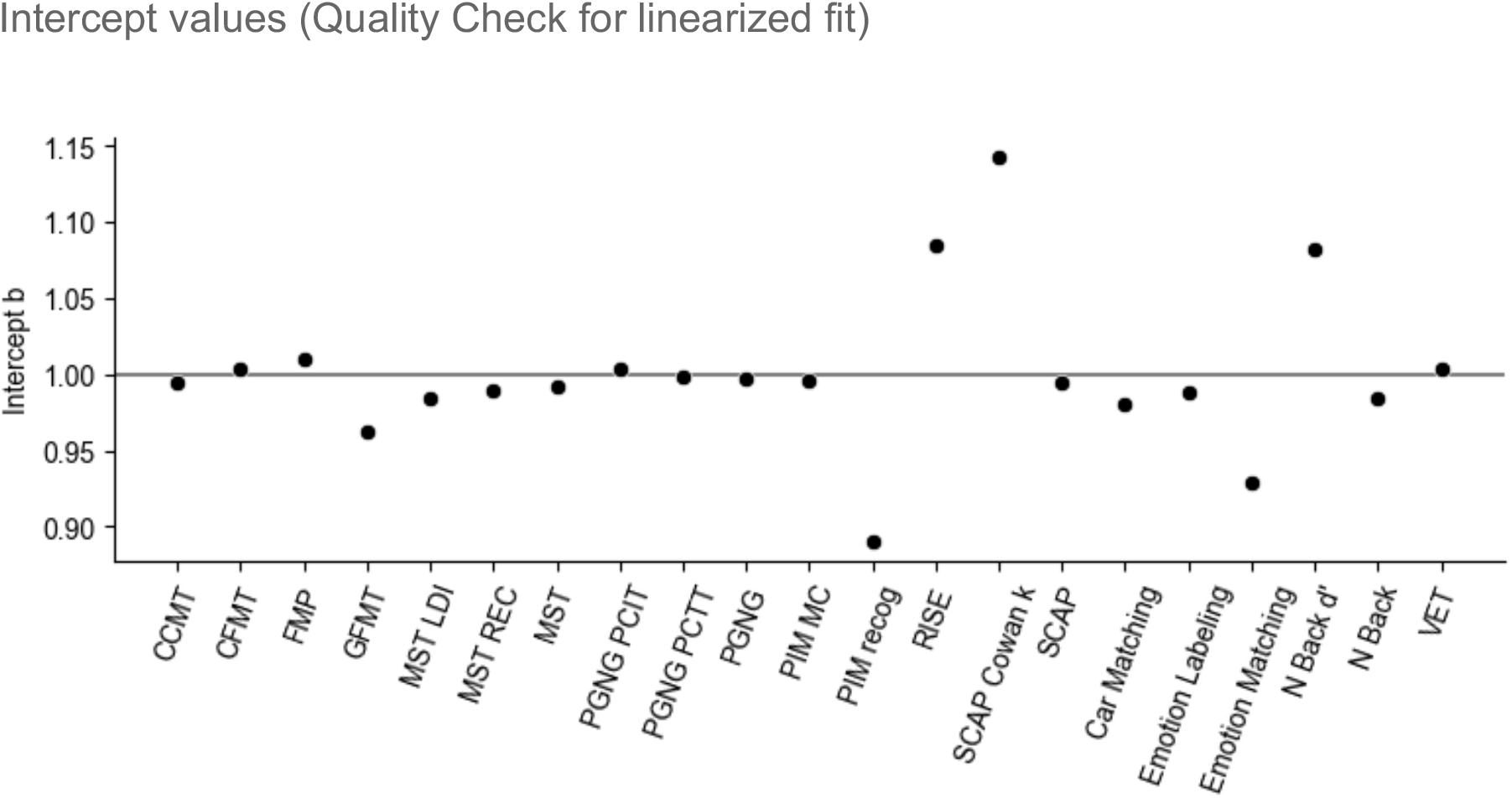
Values of intercept for linearized fit across all the measures (21). According to Eq. 4, the intercept value should be equal to one (horizontal line). Deviations of the intercept value from one suggest a violation of the assumptions (no learning / fatigue).

**Supplementary Figure 7.**
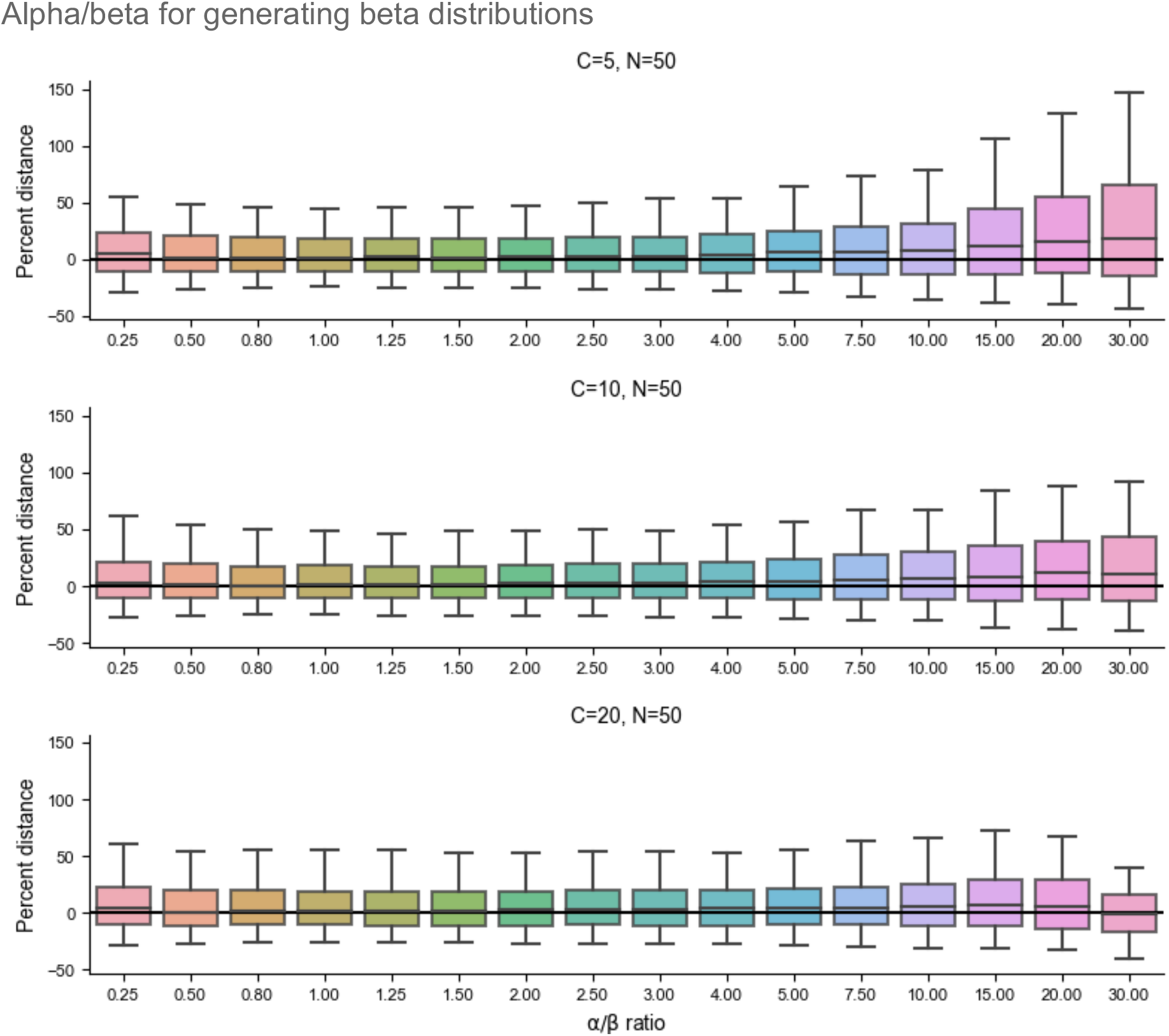
Ratios of alpha to beta (*α*/*β*) for generating beta distributions have a small effect on the percent error when estimating *C* coefficient using the MV fit.

**Supplementary Figure 8.**
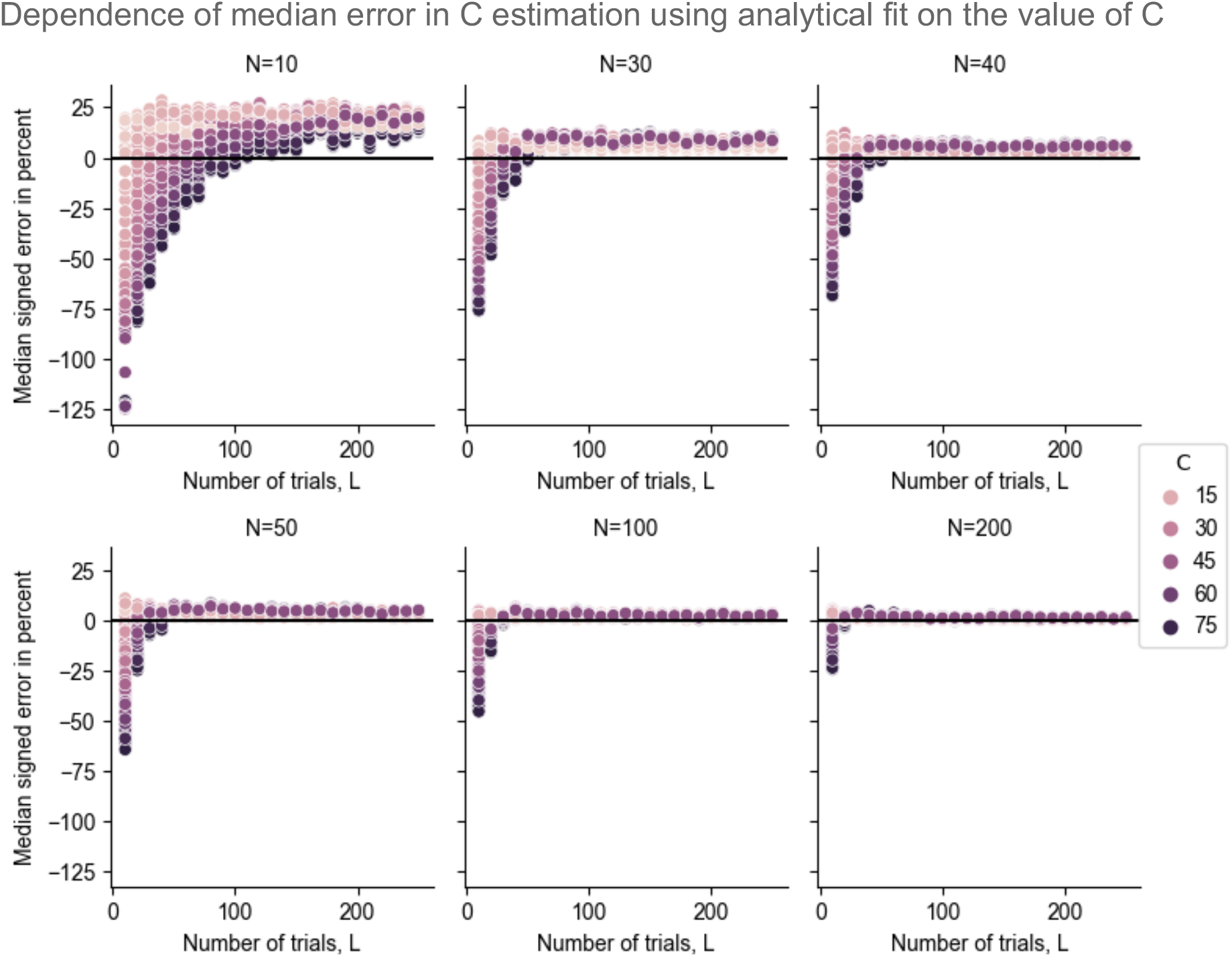
Dependence of median error in *C* estimation using the MV fit on the value of *C* for *N*=250 participants using synthetic data.

### Supplementary information

#### Instructions for use of the online tool

To use the online web application (https://jankawis.github.io/reliability-web-app/), follow this protocol:

1. Run your pilot experiment with a small number of participants and trials per participant (we recommend *L*>20, and at least *N*=30, though preferably *N*∼50. Using a smaller number of trials or participants is possible, but will result in larger confidence intervals).
2. Calculate the mean score per each participant across trials.
3. Calculate the mean (*E*[*Z*]) and sample variance (Var(*Z*)) of those scores across the population.
4. Enter the mean, variance, and the number of participants and trials per participant into the online tool and plot the corresponding reliability curve.

Toggle the reliability *R* to see how many trials *L* you need to reach your desired reliability level. You can also optionally provide time it took to collect these *L* trials and plot reliability versus the time it would take to collect the necessary number of trials.

For some distributions, for which the calculated *C* is very high, corresponding to very low variance (potentially as a result of floor or ceiling effects), the app will provide a warning. In such cases, the use of that measure for studying individual differences should be reconsidered. A warning will also be issued if the error of the fit cannot be properly estimated for very small combinations of *L* and *N*. Note that a further limitation is that the MV fit implemented in the app requires measures to be sampled from a Bernoulli distribution (i.e., each trial can yield only two possible outcomes). For any other measures which do not obey these assumptions, we suggest applying the directly fitted or linearized fits (not through the app).

#### Change in variance in constrained datasets

For the calculation of the standard deviation of the correlations between the tasks shown in Supplementary Figure 3, we first created two synthetic datasets that mirror the observed distributions of tasks from our real-world dataset. Using the mean accuracy from 75 participants who had two forms of data from each of the pairs of tasks, we generated ten synthetic forms of data per participant that preserved both the reliability of each task as well as their observed correlations. We next calculated how many trials (*L*) would be necessary for each task to achieve a given level of reliability, as was done in previous analyses. Using the ten forms of synthetic data, we created three constrained datasets, where the required number of trials *L* would comprise 80%, 50% or 20% of the total synthetic dataset. For example, if a task required 9 trials to achieve a reliability of 0.3, we would first randomly select 9/0.8 = 11 trials for our most constrained sample dataset, 9/0.5 = 18 trials for our moderately constrained sample dataset, and 9/0.2 = 45 trials for our minimally constrained sample dataset. For each level of reliability, the required *L* trials were then randomly sampled 1000 times from within these three constrained datasets. The resulting correlations (and the mean and standard deviation across the iterations) between the two pairs of tasks were therefore calculated based only on subsampling from the relevant constrained dataset. This entire simulation process was repeated 100 times, each time randomly selecting the constrained dataset from the full set of ten synthetic task forms. This yielded 100 values of standard deviations, for each reliability level, and each constrained dataset, and Supplementary Figure 3 plots the mean and standard deviation of these values.

#### Alpha/beta for generating beta distributions

To assess the impact of the ratio of the two parameters, *α* and *β*, that define the beta distribution in our simulations, we selected three values of the *C* coefficient (*C*=5, 10, 20), and for each *C*, we created 15 different combinations of *α* and *β* with the following *α*/ *β* ratios: 0.25, 0.5, 0.8, 1, 1.25, 1.5, 2, 2.5, 3, 4, 5, 7.5, 10, 15, 20, 30. Each pair of *α* and *β*, i.e., each ratio, was used to generate a beta distribution for *N*=50 participants each with *L*=250 trials. The remaining parts of the simulations were similar to the *Error estimation* of the different fits but using only the MV fit. We conducted 1000 simulations for each combination of *C* and the *α*/ *β* ratio. The ground truth for the *C* coefficient was established for every combination of *C* and *α*/*β* separately by generating a large distribution with *N* = 10^7^ and using the MV fit to get the *C* corresponding to this distribution. We computed the difference between the fitted *C* and this true *C*. We then divided this distance by the true *C* to get error in percent and plot it as boxplots in Supplementary Figure 7.

## Derivation of reliability formulas

### 1 Derivation of the closed formula

#### 1.1 Definitions and assumptions

We assume that there are *N* ≫ 1 participants performing a given task. A task consists of 2 · *L* trials, and the result of a single task is a single score per participant. We assume that each participant *i* has some value representing their “true proficiency” at the task, *p*_*i*_, and the outcome of a single trial, 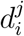, is a random variable (whose properties depend on *p*_*i*_, and with *j* an arbitrary index, since the variables 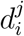 are i.i.d. random variables). We assume there is no learning and no fatigue, i.e., samples are independent. In turn, the proficiency *p*_*i*_ is also assumed to be a random variable and is assumed to be drawn independently for each participant from the continuous probability distribution *f* (*P*). Throughout, we will use *P* to denote the random variable associated with the proficiency distribution, e.g., 𝔼[*P*]= ∫ *f* (*P*)*pdp*.

We define the *sum* of the outcomes of trials performed by participant *i* in, without loss of generality, the first *L* trials as the random variable *x*_*i*_ and in the last *L* by the random variable *y*_*i*_. Since the outcomes of the trials for a participant are independent, 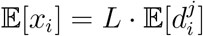 and 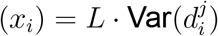.

We define two vectors of length 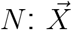, whose *i*-th element is *x*_*i*_ and 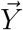, whose *i*-th element is *y*_*i*_. Our goal is to calculate the Pearson correlation between two vectors representing the average scores of participants, each across *L* trials. This is equivalent to calculating the Pearson correlation between the sum of the participants’ trial outcomes instead of their average, so we will use sums for the sake of simpler calculations.

Finally, we define a random variable *x*_*p*_, with the same properties as that of *x*_*i*_ but when fixing the proficiency to be a specific value *p*. Similarly, *d*_*p*_ will denote the performance in a single trial.

### 1.2 Calculation

The Pearson correlation coefficient is defined by:

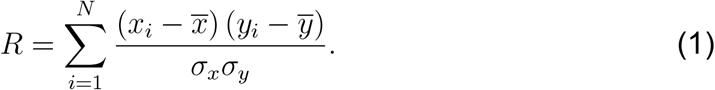

where 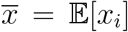 and 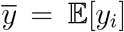 denote the means of 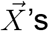 and 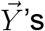 elements respectively, and *σ* their corresponding standard deviations.

For the next step, we group the participants by proficiency. The inner sum is over all the participants with a given proficiency 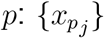

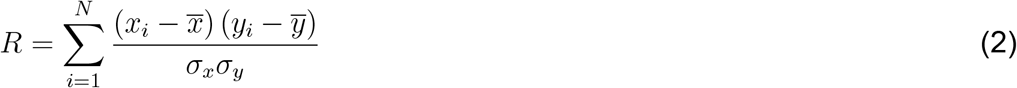

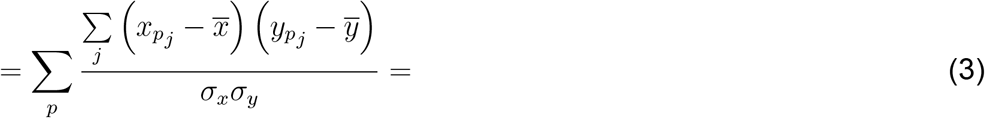

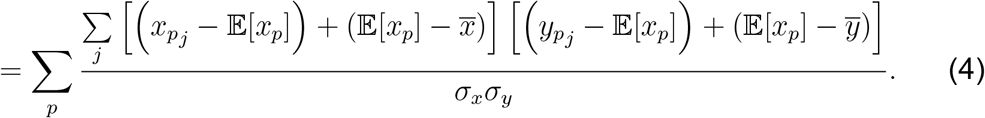

It is clear that 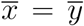 and *σ*_*x*_ = *σ*_*y*_. Using those facts and substituting 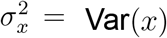, we can further simplify the equation into the following formula:

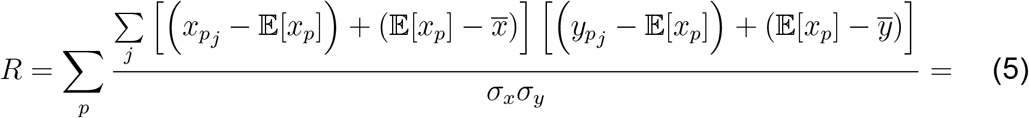

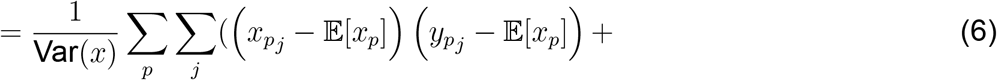

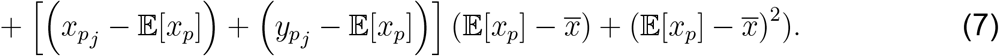

Since 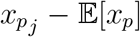 and 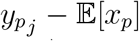 are fluctuations around their respective expected values, and 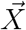 and 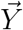 are independent, the averages of their products vanish. Therefore, we obtain:

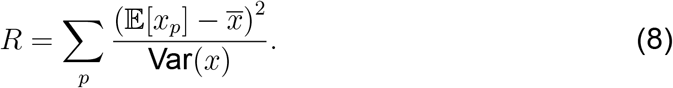

Next, we calculate 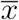:

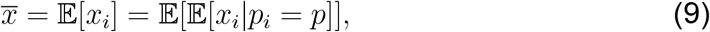

with 𝔼[*x*_*i*_ | *p*_*i*_ = *p*] = 𝔼[*x*_*p*_] denoting conditioning on the proficiency of the participant being equal to *p*, and the outer expectation value is the average over all participants (accounting for the distribution of proficiencies). This leads to:

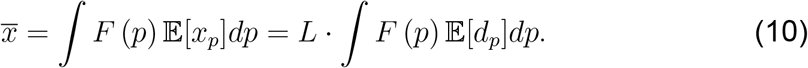

Additionally:

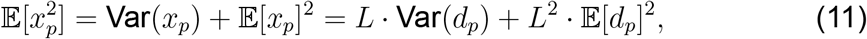

and:

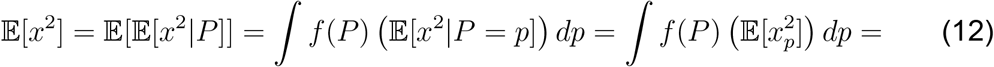

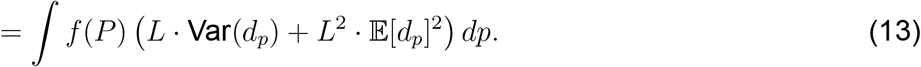

Using that, we can calculate Var(*x*):

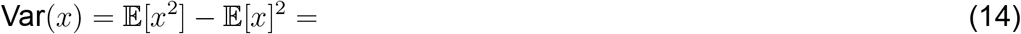

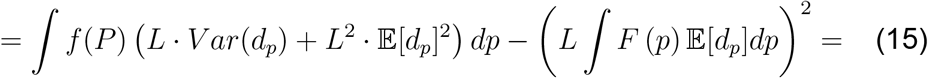

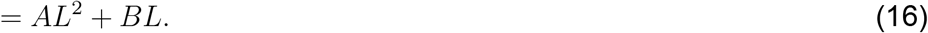

where we defined the two new coefficients (independent of *L*):

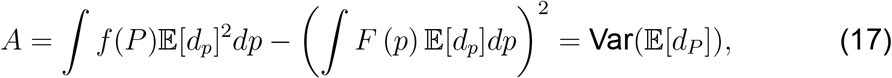

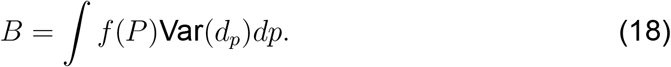

Returning to Eq. 8 and using what we derived, we can express reliability in the following way:

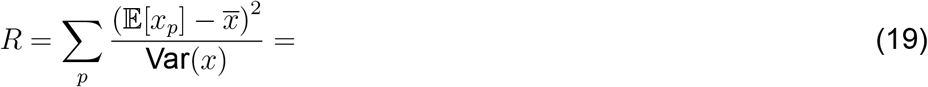

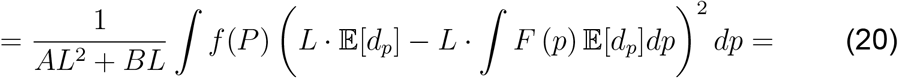

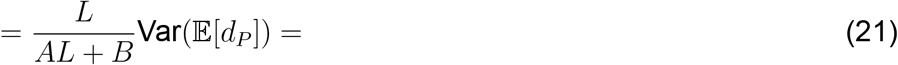

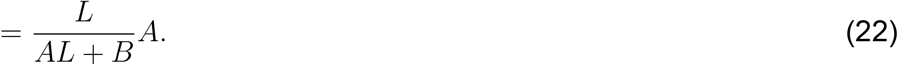

That leads us to the final formula:

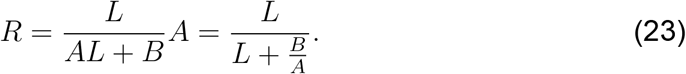

We can denote by *C* the ratio *B*/*A*:

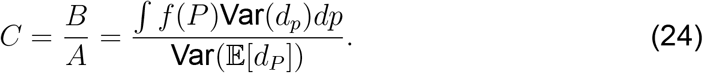

The relationship between *L* and *R* that we derived can be expressed as

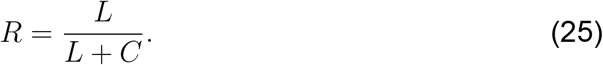

Note that this is equivalent to the famous Spearman-Brown formula.^1,2^

## 2 Particular Case – Tasks with Binary outcomes

If the tasks given to the participants have binary outcomes (e.g., 0/1, correct/incorrect, …), we have *d*_*p*_ ∼ *Bernoulli*(*p*). Using that, we can derive an explicit formula for *R*(*L*). Following from that, we know:

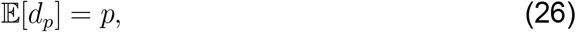

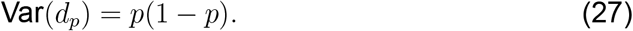

We can calculate *C* explicitly:

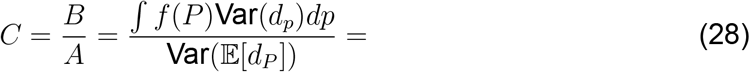

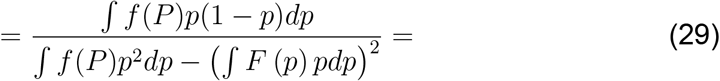

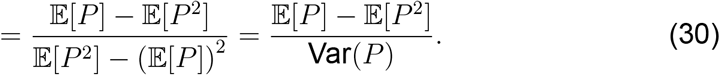

We know that:

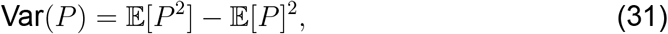

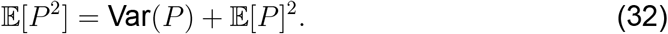

So we can simplify the expression into:

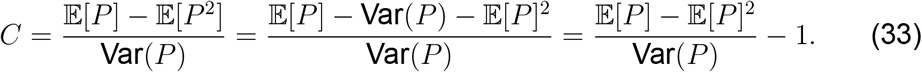

### 2.1 Calculating C for the beta distribution

For the beta distribution defined by two parameters, *α* and *β*, we can derive the value of C explicitly. The mean and the variance of beta distribution can be expressed using those two parameters in the following way:

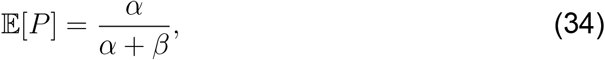

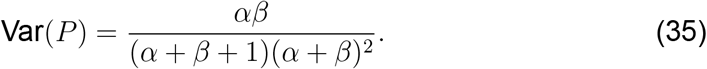

Substituting to Eq. 33, we get:

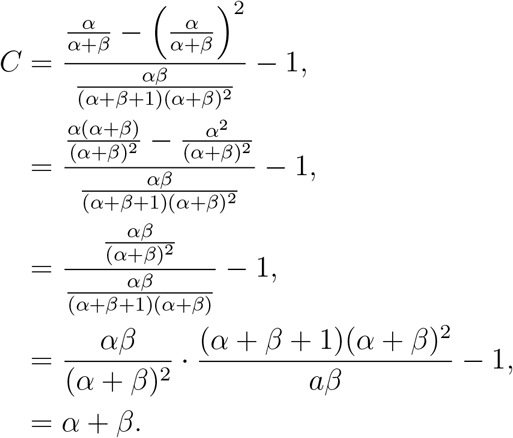

## 3 Computing Var(P) and 𝔼[*P*^2^] from the participants’ scores

### 3.1 Background and assumptions

The previous calculations derived the Pearson correlation coefficient, theoretically, for a population size → ∞ (i.e., this is implicit when we take the ensemble average). When handling real data, we can only calculate the Pearson correlation coefficient for a finite population size. This correlation coefficient would approach the theoretical value as *N* → ∞. Since we use a finite number of participants, we should expect some error between the theoretical value of the coefficient and the value calculated from a finite population, but no bias.

There is, however, another source of error. When trying to calculate *C* from real data, we never have access to the real participants’ proficiencies. We can never know *P*, and we are thus introducing an error by having a limited number of trials per participant when estimating their proficiencies.

We have, nonetheless, the scores of the participants (*x*_*i*_ for the *i*-th participant) from which we can estimate the *p*_*i*_ of the *i*-th participant using 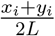. This naive approach, however, creates a bias when calculating 𝔼[*P*^2^] and Var(*P*). As these are used in Eq. 33, we would expect to see a bias if we used this equation in this form (as shown in Supplementary Fig. 4). In this section, we derive a formula that does not have this bias and will allow us to estimate *C* from data accurately.

For simplicity, we assume a binary case, where a participant’s score with proficiency *p* will be given by a random variable with distribution *x*_*p*_ ∼ *B*(2*L, p*) (binomial). We define a random variable 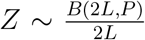 that provides an estimate of the participant’s proficiency.

Our goal in this section is to calculate how Var(*Z*) depends both on Var(*P*) and on *L*, enabling us to express *C* using the statistics of *Z* and thus overcoming the bias stemming from calculating the proficiencies naively when given a limited number of trials per participant.

### 3.2 Calculating Var(P)

The law of total variance dictates that:

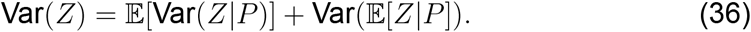

Using:

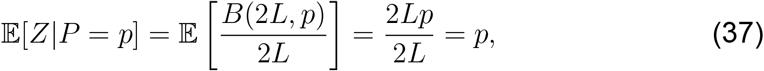

we can calculate:

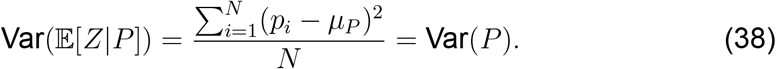

Next, we use:

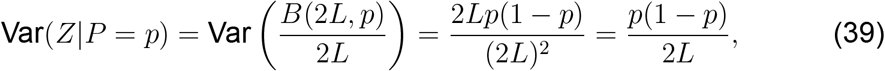

to calculate:

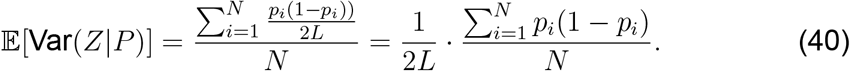

Putting it together, we find:

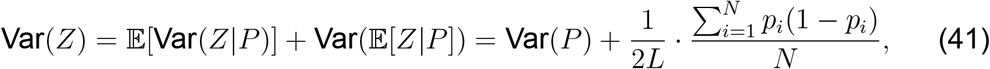

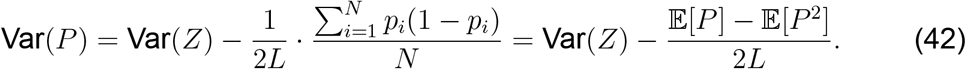

#### 3.3 **Calculating** 𝔼[*P* ^2^]

We observe that:

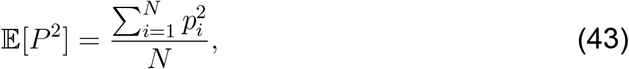

and:

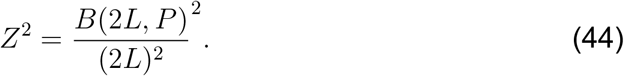

From the expected value of the square of a binomial variable, we can now see that:

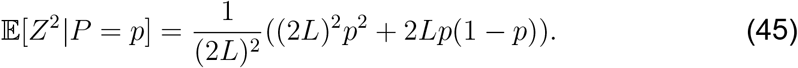

Using the law of total expectation, we can now calculate the following:

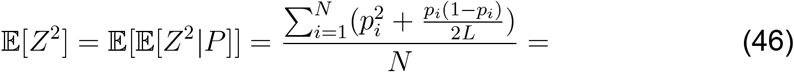

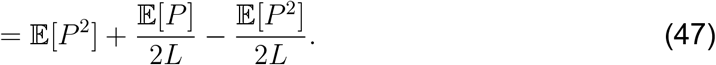

Similarly,

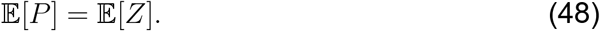

Finally, we obtain:

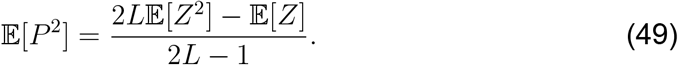

### 3.4 Calculating C

We know that:

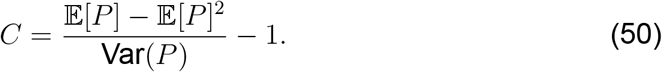

Using Eqs. 42 and 49, we can derive:

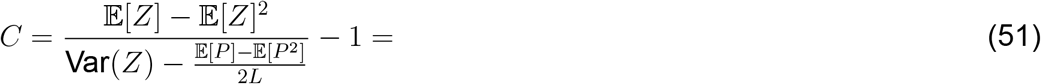

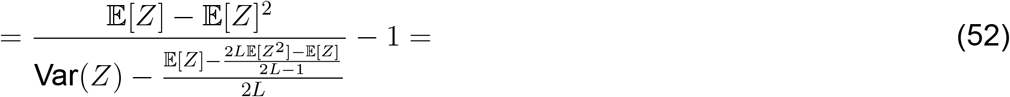

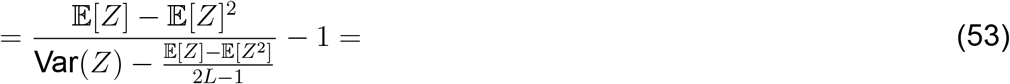

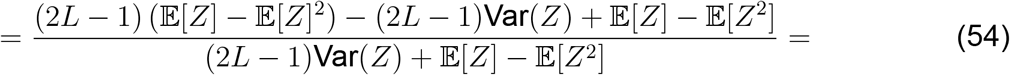

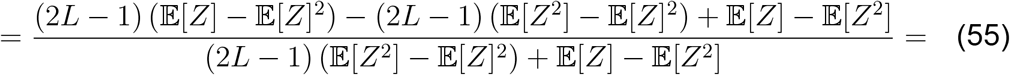

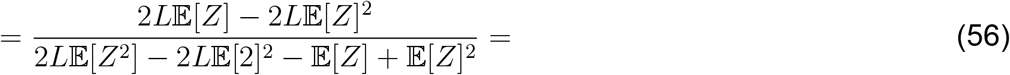

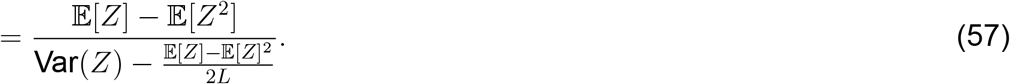

Note that this is an explicit formula for the coefficient *C* using *observed* data alone. Eq. 57 is reminiscent of an equation previously described by Kuder and Richardson (KR-21^3–5^).

## 4 Derivation of the Attenuation-Correction Formula

In this section, we derive the Attenuation-Correction formula, first formulated by Spearman^1,6^ and further developed by others,^7–9^ to represent the correlation between two quantities represented by vectors of their measurements, 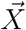 and 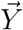, using the internal reliabilities of those measurements. We start by defining 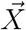 and 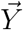 as:

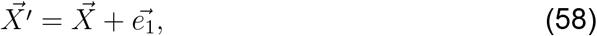

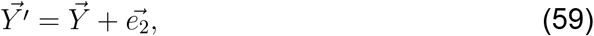

where 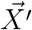, 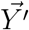 are vectors representing the measured scores, 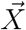, 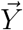 are vectors representing the real, underlying proficiencies, and 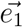, 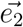 are the errors from the measurement, respectively. We assume that 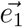, 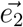 are independent, and also that 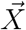, 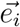 are independent (same for 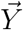). We note that the Pearson correlation between two variables can be written as:

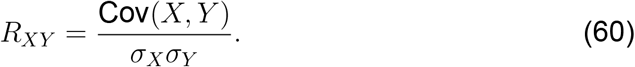

If we denote 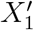 and 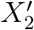 as the split-halves of *X*^*′*^, we can use Eq. 60 to see that the internal reliabilities of the measurement vectors are:

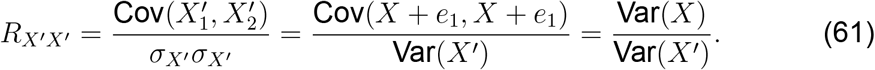

Analogous formula gives *R*_*Y′Y′*_. Now we can calculate the following:

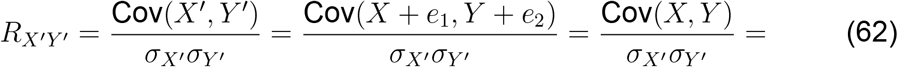

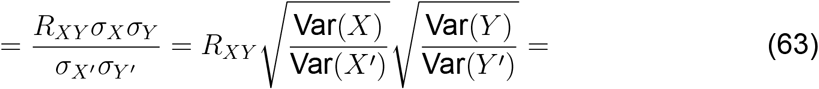

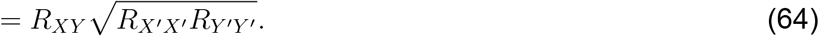

